# Dopamine supports reward prediction to shape reward-pursuit strategy

**DOI:** 10.1101/2025.08.22.671841

**Authors:** Melissa Malvaez, Andrea Suarez, Nicholas K. Griffin, Kathia Ramírez-Armenta, Sean B. Ostlund, Kate M. Wassum

**Affiliations:** Dept. of Psychology, UCLA, Los Angeles, CA 90095, USA; Department of Anesthesiology. UCI, Irvine, CA; Brain Research Institute, UCLA, Los Angeles, CA 92697, USA

**Keywords:** Pavlovian conditioning, Pavlovian-to-instrumental transfer, Nucleus accumbens, Ventral tegmental area, Fiber photometry, Optogenetics

## Abstract

Reward predictions not only promote reward pursuit, they also shape how reward is pursed. Such predictions are supported by environmental cues that signal reward availability and probability. Such cues trigger dopamine release in the nucleus accumbens core (NAc). Thus, here we used dopamine sensor fiber photometry, cell-type and pathway-specific optogenetic inhibition, Pavlovian cue–reward conditioning, and test of cue-induced reward-pursuit strategy in male and female rats, to ask whether cue-evoked phasic dopamine release is shaped by reward prediction to support reward pursuit. We found that cue-evoked NAc core dopamine is positively shaped by reward prediction and inversely relates to and predicts instrumental reward seeking. Cues that predicted imminent reward with high probability triggered a large NAc dopamine response and this was associated checking for the expected reward in the delivery location, rather than instrumental reward seeking. Cues that predicted reward with low probability elicited less dopamine and this was associated with a bias towards seeking, rather than check for reward. Correspondingly, inhibition of cue-evoked NAc dopamine increased instrumental reward-seeking and decreased reward-checking behavior. Thus, transient, cue-evoked NAc core dopamine release supports reward prediction to shape reward-pursuit strategy.

**SIGNIFICANCE STATEMENT:** Cues that signal reward availability can cause us to pursue reward. To ensure this is adaptive, we use the predictions these cues enable to help us decide how to pursue reward. When reward prediction is low, we’ll seek out new reward opportunities. When it is high, we’ll exploit the expected reward by checking for it in its usual location. Here we discovered that transient, cue-evoked nucleus accumbens dopamine fluctuations support reward predictions to shape how reward is pursued. The data show that dopamine can actually constrain reward seeking and promoting reward checking when reward is predicted strongly and imminently. These results provide new information on how dopamine shapes behavior in the moment and help understand the link between motivational and dopamine disruptions in psychiatric conditions such as addictions and depression.

---

Reward cues can heavily influence our behavior. By signaling reward availability and probability, they support reward predictions that not only promote reward pursuit (Corbit and Balleine, 2016; Robinson and Berridge, 2025), but also influence *how* we pursue reward (Ostlund and Marshall, 2021). When a cue only sporadically signals a desired resource, it is adaptive to engage in instrumental ‘reward-seeking’ actions. But if reward is more reliably predicted, it is better to pursue it by ‘checking’ its usual delivery location (Azrin and Hake, 1969; Lovibond, 1981; Stephens and Krebs, 1986; Timberlake, 1994; Anselme and Güntürkün, 2018; Marshall and Ostlund, 2018). This is particularly true for cues that signal imminent reward. When time is of the essence, seeking reward can prevent retrieval of an expected reward, and vice versa. So, we tend to seek reward when its presence is unlikely, but check for it when it is probable. For example, with only a few minutes before a meeting, if snacks are offered only sporadically, you might seek one out at the vending machine, but, if they are more reliably predicted, you’ll head straight to the conference room to look for a treat. Thus, reward prediction is critical for adaptive behavior. Inability to adaptively regulate reward pursuit can lead to unnecessary opportunity costs, loss of control, or maladaptive behaviors. This can underlie symptoms of psychiatric conditions such as addictions and depression (Levy and Dubois, 2006; Jentsch and Pennington, 2014; Le Heron et al., 2018; Robinson and Berridge, 2025). Yet, little is known of how the brain supports reward prediction to shape reward-pursuit strategy.

Dopamine may contribute. Reward-predictive cues trigger dopamine release in the nucleus accumbens core (NAc) (Roitman et al., 2004; Day et al., 2007; Day et al., 2010; Wassum et al., 2013; Aitken et al., 2015; Saddoris et al., 2015; Collins et al., 2016; Sun et al., 2018; Kutlu et al., 2021; Garr et al., 2024; Bornhoft et al., 2025). In motivational accounts, this phasic signal is a critical substrate through which cues promote reward-pursuit behaviors (Beeler et al., 2012; Saunders and Robinson, 2012; Wassum et al., 2013; Syed et al., 2016; Berke, 2018; Robinson and Berridge, 2025). Such cue-evoked dopamine can be shaped by elements of reward prediction (Gan et al., 2009; Day et al., 2010; Hamid et al., 2015; Hart et al., 2015; Saddoris et al., 2015; Aitken et al., 2016; Garr et al., 2024). This raises the possibility that cue-evoked NAc dopamine may support reward prediction to shape how reward is pursued. Thus, here we investigated the relationship between cue-evoked NAc dopamine and reward-pursuit behaviors and asked whether dopamine supports reward prediction to shape the strategy with which reward is pursued. To achieve this, we combined NAc dopamine measurement and manipulation with Pavlovian cue–reward conditioning and a Pavlovian-to-instrumental transfer test of the influence of cues signaling different reward probabilities on reward-pursuit strategy.

## METHODS

### Subjects

Subjects were male and female wildtype Long-Evans rats and transgenic Long-Evans rats expressing Cre recombinase under control of the tyrosine hydroxylase promoter (Th-cre+) aged 10 - 12 weeks at the time of surgery. Rats were housed in a temperature (68 - 79°F) and humidity (30 - 70%) regulated vivarium. They were initially housed in same-sex pairs and then, following surgery, housed individually to preserve implants. Rats were provided with water *ad libitum* in the home cage and, during experimental training and testing, were maintained on a food-restricted 12 - 18-g daily diet (Lab Diet, St. Louis, MO) to maintain approximately 85 - 90% free-feeding body weight. Rats were habituated to handling for 3 - 5 days prior to the onset of each experiment. Separate groups of naïve rats were used for each experiment. Experiments were performed during the dark phase of a 12:12 hr reverse dark:light cycle (lights off at 7 AM). All procedures were conducted in accordance with the NIH Guide for the Care and Use of Laboratory Animals and were approved by the UCLA Institutional Animal Care and Use Committee.

### Surgery

Rats were anesthetized with isoflurane (4 - 5% induction, 1 - 2% maintenance) and positioned in a digital stereotaxic frame (Kopf, Tujunga, CA). Subcutaneous Rimadyl (Carprofen; 5 mg/kg; Zoetis, Parsippany, NJ) was given pre-operatively for analgesia and anti-inflammatory purposes. An incision was made along the midline to expose the skull. After performing a small craniotomy, virus was injected using a 28-g infusion needle (PlasticsOne, Roanoke, VA) connected to a 1-mL syringe (Hamilton Company, Reno, NV) by intramedic polyethylene tubing (BD; Franklin Lakes, NJ) and controlled by a syringe pump (Harvard Apparatus, Holliston, MA). Virus was injected at a rate of 0.1 µl/min and the needle was left in place for 10 min post-injection to allow diffusion. Further experiment-specific surgical details are provided below. After surgery, rats were kept on a heating pad maintained at 35 °C for 1 hr and then single housed in a clean homecage for recovery and monitoring. Rats received chow containing the antibiotic Trimethoprim/Sulfamethoxazole (TMS; Inotiv, West Lafayette, IN) for 7 days following surgery to prevent infection, after which they were returned to standard rodent chow.

### Behavioral procedures

#### Apparatus

Training took place in Med Associates conditioning chambers (East Fairfield, VT) housed within sound- and light-attenuating boxes, described previously (Malvaez et al., 2015; Collins et al., 2019). Each chamber had grid floors and contained a retractable lever that could be inserted to the left of a recessed food-delivery port (magazine) on the front wall. A photobeam entry detector was positioned at the entry to the food port. Each chamber was equipped with a pellet dispenser to deliver 45-mg food pellets (Bio-Serv, Frenchtown, NJ) into a well of the food port. A tone generator and clicker were attached to a speaker on the wall opposite the lever and food-delivery port. A fan mounted to the outer chamber provided ventilation and external noise reduction. A 3-watt, 24-volt house light mounted on the top of the back wall opposite the food port provided illumination. For optogenetic manipulations, chambers were outfitted with an Intensity Division Fiberoptic Rotary Joint (Doric Lenses, Quebec, QC, Canada) connecting the output fiber optic patch cords to a laser (Changchun New Industries Optoelectronics Technology Co., ChangChun, JiLin, China) positioned outside of the chamber.

#### Magazine conditioning

Rats first received 2 sessions (1 session/day) of training to learn where to receive the food-pellet reward (45-mg grain pellet; Bio-Serv). Each session included 30 non-contingent pellet deliveries (60-s intertrial interval, ITI).

#### Short-delay Pavlovian conditioning

All rats received 8 sessions of short-delay Pavlovian conditioning (1 session/day) to learn to associate each of 2, 10-s auditory (75 - 78 db) cues (aka conditioned stimuli, CS), click (10 Hz) and tone (1.5kHz), with the delivery of grain pellets. For 10 subjects in the fiber photometry experiment the tone was continuous for 10-s, for all other subjects, the tone was pulsed in 0.25-s on/off intervals (4 Hz). Each 10-s cue co-terminated with the delivery of a single pellet. The high-probability cue (CS_90_) co-terminated with reward delivery on 90% of the trials. The low-probability cue (CS_30_) co-terminated with reward delivery on only 30% of the trials. For half the subjects, click served as the CS_90_ and tone as the CS_30_, with the other half receiving the opposite arrangement. Each session consisted of 20 click and 20 tone presentations. Cues were delivered pseudo-randomly with a variable 60 - 160-s ITI (mean = 110 s).

#### Instrumental conditioning

Rats next received 8 sessions, minimum, of instrumental conditioning (1 session/day), in which lever-press actions were reinforced with delivery of a single grain pellet. Each session terminated after 30 outcomes had been earned or 40 min had elapsed. Lever presses were continuously reinforced during the first session and then the reinforcement schedule was shifted to random interval (RI) on which a variable interval must elapse following a reinforcer for another press to be reinforced. The variable interval started at an average of 15 s (RI-15s) for one session, then escalated to 30 s (RI-30s) for one session, and finally averaged 60 s (RI-60s) for the remaining sessions.

#### Pavlovian retraining

Following instrumental training, rats received 4 additional sessions of Pavlovian conditioning, as described above.

#### Pavlovian-to-instrumental transfer test

Rats next received a Pavlovian-to-instrumental transfer (PIT) test. During the PIT test, the lever was continuously present, but pressing was not reinforced. After 10 min of lever-pressing extinction, each 10-s cue was presented separately 10 times, separated by a fixed 110-s ITI. Cues were presented in pseudorandom order with no more than 2 of the same cue type occurring in succession. The cue type that started the session was counterbalanced across subjects. No rewards were delivered following cue presentation.

#### Behavioral data collection and analysis

Entries into the food-delivery port and/or lever presses were recorded continuously for each session and processed with Microsoft Excel (Microsoft, Redmond, WA). For Pavlovian conditioning, conditional food-port approach responses were assessed by comparing entries into the food-delivery port during the 10-s cue to the 10-s baseline periods immediately prior to cue onset (precue). Data were averaged across trials and then averaged across the last 2 conditioning sessions. For instrumental conditioning, press rates (presses/min) were averaged across the last 2 training sessions. For PIT tests, we evaluated entries into the food-delivery port and lever presses for a 15-s cue window combining the 10-s cue and 5-s post-cue period. This allowed us to capture behavior during the cue itself and the post-cue period, when reward was delivered during training, and, thereby capture the delayed excitatory effect of short-delay cues and distinguish the effect of strongly v. weakly predictive cues (Marshall et al., 2020). See Extended Data Figures 1-1 and 2-1 for a breakdown of food-port entries and lever presses in 2-s bins before, during, and after cue presentation during PIT. Lever presses served as the measure of instrumental reward seeking. Presses during the 15-s cue periods were compared to that during the 15-s precue baseline. As shown previously (Halbout et al., 2019; Halbout et al., 2022), we found that many food-port entries occurred immediately (within 2 s) following a lever press (Extended Data Figure 1-2). Therefore, to isolate Pavlovian conditional food-port checks from these entries that occurred as part of an instrumental press-check sequence, we excluded food-port entries that directly followed a lever press within 2 s. The remaining food-port entries were our measure of reward-checking behavior. All food-port entries and press-linked food-port entries are shown in Extended Data Figure 1-2 and 2-2. Food-port entries during the 15-s cue periods were compared to that during the 15-s precue baseline. To account for baseline behavior rates and evaluate the cue-induced change in behavior, we also computed the change (Δ) in lever presses and unlinked food-port entries by subtracting those during the 15-s cue period from baseline. Data were averaged across trials for each cue type. Trial x trial results are presented in Extended Data Figure 1-4.

### Fiber photometry recordings of NAc dopamine release during Pavlovian-to-instrumental transfer

#### Subjects

Subjects were naïve, male and female Long Evans rats (Final *N* = 14, 7 male). 2 subjects were removed from the experiment: 1 due to illness and 1 because it did not acquire Pavlovian conditional behavior. 3 subjects with tissue damage that prevented placement validation were excluded from the analysis.

#### Behavioral training

Rats were food restricted and received magazine conditioning, followed by Pavlovian and instrumental training as above. Following surgery (see below) and recovery, rats were again food restricted and received 4 additional Pavlovian and 4 additional instrumental training sessions. Rats were tethered for the retraining sessions, but no light was delivered.

#### Surgery

Rats were infused unilaterally (Right hemisphere *N* = 13, Left: *N* = 1) with AAV encoding the GPCR-activation-based dopamine sensor GRAB_DA2m_ (pAAV9-hsyn-GRAB_DA4.4, Addgene, Watertown, MA). Virus (0.5 µl) was infused at 0.1 µl/min into the NAc (AP: +1.3; ML: ±1.3; DV: −7.2, from bregma). 5 min later, viral injectors were dorsally repositioned in the NAc for a second viral infusion (0.5 µl; DV: −6.4). Optical fibers (200-µm diameter, 0.37 NA, Neurophotometrics Ltd, San Diego, CA) were implanted (DV: −6.8). Rats were allowed to recover for at least 5 days before retraining commenced.

#### Fiber photometry recordings during Pavlovian-to-instrumental transfer

Fiber photometry was used to image GRAB_DA2m_ fluorescent changes in NAc neurons during PIT using a commercial fiber-photometry system (Neurophotometrics Ltd., San Diego, CA). 470-nm excitation light was adjusted to approximately 80 - 100 µW at the tip of the patch cord (fiber core diameter: 400 µm; Doric Lenses, Quebec City, Canada). Fluorescence emission was passed through a 535-nm bandpass filter and focused onto the complementary metal-oxide semiconductor (CMOS) camera sensor through a tube lens. Samples were collected at 20 Hz using a custom Bonsai (Lopes et al., 2015) workflow. Time stamps of task events were collected simultaneously through an additional synchronized camera aimed at the Med Associates interface, which sent light pulses coincident with task events. Signals were saved using Bonsai software and exported to MATLAB (MathWorks, Natick, MA) for analysis.

#### Fiber photometry analysis

Data were pre-processed using a custom-written pipeline in MATLAB (MathWorks, Natick, MA)(Sias et al., 2021; Sias et al., 2024). The 415-nm and 470-nm signals were fit using an exponential curve. Change in fluorescence (ΔF/F) at each time point was calculated by subtracting the fitted 415-nm signal from the 470-nm signal and normalizing to the fitted 415-nm data [(470-fitted 415)/fitted 415)]. Non-normalized 470-nm signal is provided in Extended Data Figure 1-5. The ΔF/F data was Z-scored to the average of the whole session [(ΔF/F - mean ΔF/F)/std(ΔF/F)]. Z-scored traces were then aligned to behavioral event timestamps. Individual trial data were excluded if the patch cord became detached or if artifactual signal due to excessive motion or patch cord twisting was detected (n = 6 CS_30_ trials from 4 subjects; n = 7 CS_90_ trials from 6 subjects).

Using a waveform analysis (Jean-Richard-Dit-Bressel et al., 2020), event-related activity was analyzed 5 s before to 20 s after cue onset. To ensure independence of observations, the analysis was performed at the subject-level average waveform. For each subject, the CS_30_ and CS_90_ trials were independently averaged to generate a mean time-series for each CS type for each subject. The observed difference waveform (mean CS_30_ – mean CS_90_) was analyzed for statistical significance at every time point using two non-parametric approaches: a two-sample permutation test and a bootstrap confidence interval. For the permutation test, with 1000 iterations a null distribution of condition differences was generated by randomly assigning the CS condition labels and calculating the difference of the resampled means across the entire time window. The *P* value at each time point was the proportion of permutations whose mean difference values were more extreme than the observed difference between the actual samples. A time point was flagged as significant if *P* < 0.05. To provide a robust estimate of the true population mean difference, a matrix of 1000 bootstrapped means were generated by resampling with replacement the subject-level average waveform for both CS_30_ and CS_90_. The difference between CSs was calculated for each of the 1000 iterations. CI for each timepoint were percentiles at that timepoint of the bootstrap matrix (95%: 2.5, 97.5 percentiles), which were then expanded by a factor of √n/(n−1) to counter small sample narrowness bias. A significant difference was flagged whenever the CI did not contain the null of 0. For both approaches, a threshold of 10 consecutive time points was applied to the initial set of significant indices to correct for isolated, random fluctuations. Using this approach, we found that the cue-evoked dopamine differed between cue-type immediately after CS onset. Thus, to further quantify dopamine fluctuations, we calculated the peak value in the 2-s period immediately following cue onset and compared to the 2-s precue baseline periods. To quantify dips in dopamine detected following cue offset, when the predicted reward was omitted at test, we calculated the minimum GRAB_DA_ Z-score in the 5-s postcue period compared to 5-s period before cue offset. To quantify dopamine related to entry and lever-press behaviors during the cue, we used the 4-s period prior to entries or lever presses compared to the baseline 4-s precue periods. Quantifications and signal aligned to events were averaged across trials within a session and compared between groups.

Because lever pressing and entries into the food-delivery port can freely occur during cue presentation, we used a linear regression model to parse the extent to which dopamine fluctuations are explained by these discrete, but overlapping events (Parker et al., 2016; Greenstreet et al., 2025). To calculate kernels that correspond to the isolated response to each behavioral event for each CS type: CS onset, CS offset, lever presses, and food-port entries, the Z-scored GRAB_DA_ signal 5 s prior to and 20 s after cue onset was modeled as the sum of the response to each event. Discrete events cue onset, offset, lever press, and food-port entry were convolved into a time series representing the time of the event in a series of values between 0 – 1 using a 4-s gaussian distribution. For CS onset and offset, we used a half-gaussian convolution (0 s – 4 s). Because behavioral events include both movements to make the action and a following outcome (e.g., non-reinforcement), the press and food-port entry convolution was centered on the event (−2 – 2 s). For each CS type for each subject, the coefficients of the kernels were solved using the method of least-squares with the MATLAB function *regress*. The kernel output for the 5 s prior to and 20 s after cue onset was averaged across trials for each CS type for each subject. To calculate the percentage of variance explained by each regressor class, the predicted dopamine signal was recalculated removing one regressor class (e.g., CS, both CS_30_ and CS_90_) at a time. The percent variance explained by the removed regressor was calculated by subtracting the variance explained by the model with the regressor removed from the explained variance of the full model, and expressing this as a percentage of the variance explained by the full model [(*v*_full_ - *v*_partial_/*v*_full_)*100).

### Optogenetic inhibition of VTA_DA_→NAc terminals at cue onset during Pavlovian-to-instrumental transfer

#### Subjects

Subjects were naïve, male and female Th-cre+ (hemizygous) Long Evans rats and wildtype (Th-cre-) littermates (Control, Final *N* = 12, 6 male, 9 Th-cre+/cre-dependent tdTomato, 3 Th-cre-/cre-dependent ArchT; ArchT, *N* = 14 Th-cre+, 9 male). Subjects with misplaced optic fibers (ArchT, *N* =1), lack of terminal expression (*N* = 1), or broken optic fibers (ArchT, *N* = 4; Control, *N* = 1) were excluded from the dataset.

#### Surgery

Th-cre+ rats were randomly assigned to a viral group and received bilateral infusions (0.2 µl) of AAV encoding a cre-inducible inhibitory opsin archaerhodopsin T (AAV5-CAG-FLEX-ArchT-tdTomato, Addgene), or a tdTomato fluorescent protein control (tdTomato; AAV5-CAG-FLEX-tdTomato, University of North Carolina Vector Core, Chapel Hill, NC) into the VTA (AP: −5.3; ML: ±0.7; DV: −8.3 mm from bregma). Th-cre- rats all received bilateral infusions (0.2 µl) of AAV encoding an mCherry fluorescent protein control (AAV8-hSyn-mCherry). Optical fibers (200-µm core, 0.39 NA, Thorlabs, Newton, NJ) held in ceramic ferrules (Kientec Systems, Stuart, FL) were implanted bilaterally in the NAc (AP: +1.3; ML: ±1.4; DV: −6.8, from bregma). Rats were allowed to recover for 3 - 4 weeks prior to the start of food restriction and training.

#### Behavioral training

Rats received magazine, Pavlovian, and instrumental conditioning as above. They were attached to the optical tether (200 µm, 0.22 NA, Doric Lenses) throughout Pavlovian and instrumental conditioning, but no light was delivered. One cohort (*N* = 6, 3 control, 3 ArchT) received 12 rather than 8 Pavlovian conditioning sessions.

#### Optogenetic inhibition of VTA_DA_→NAc projections during the Pavlovian-to-instrumental transfer test

The PIT test occurred 7 - 9 weeks post-surgery to allow sufficient ArchT expression in VTA_DA_→NAc terminals at the time of manipulation. Optogenetic inhibition was used to attenuate the activity of ArchT-expressing VTA_DA_ axons and terminals in the NAc at cue onset. Green light (532 nm; 10 mW) was delivered to the NAc via a laser (Changchun New Industries Optoelectronics Technology Co.) connected through a ceramic mating sleeve (Thorlabs) to the ferrule implanted on the rat. During the PIT test, 2.5 s of continuous light was delivered beginning 0.5 s before each cue onset. Light effects were estimated to be restricted to the NAc core based on predicted irradiance values (https://web.stanford.edu/group/dlab/cgi-bin/graph/chart.php). Individual trials in which the patch cord became detached, and, therefore, light was not delivered, were removed from the analysis (Controls: n = 1 CS_90_ trial in 1 subject; ArchT: n = 3 CS_30_ trials across 3 subjects; n = 5 CS_90_ trials across 4 subjects). Because behavioral data did not differ between the Th-cre+ and TH-cre- control types (PIT lever presses: 3-way RM-ANOVA, Control type: F_1,20_ = 1.52, *P* = 0.23; Control type x CS presence: F_1,20_ = 2.11, *P* = 0.16; Control type x CS type: F_1,20_ = 0.13, *P* = 0.72; Control type x CS presence x CS type: F_1,20_ = 0.67, *P* = 0.42; PIT food-port entries: 3-way RM-ANOVA, Control type: F_1,20_ = 0.01, *P* = 0.94; Control type x CS presence: F_1,20_ = 0.02, *P* = 0.89; Control type x CS type: F_1,20_ = 0.22, *P* = 0.64; Control type x CS presence x CS type: F_1,20_ = 1.80, *P* = 0.19), the controls were collapsed into a single group.

### Histology

Following behavioral experiments, rats were deeply anesthetized with pentobarbital (Vortech Pharmaceuticals, Dearborn, MI) and transcardially perfused with phosphate buffered saline (PBS) followed by 4% paraformaldehyde (PFA). Brains were removed and post-fixed in 4% PFA overnight, placed into 30% sucrose solution, then sectioned into 30-μm slices using a cryostat (CM1950, Leica Biosystems, Nusslock, Germany) and stored in cryoprotectant. Slices were rinsed in a DAPI (Thermofisher Scientific, Waltham, MA) solution for 4 min (5 mg/mL stock, 1:10000), washed 3 times in PBS for 15 min, mounted on slides and coverslipped with ProLong Gold mounting medium (Thermofisher Scientific). All images were acquired using a fluorescence microscope (BZ-X710; Keyence, Osaka, Japan) with 4X, 10X, and 20X objectives (CFI Plan Apo), CCD camera, and BZ-X Analyze software (Keyence), to confirm viral expressions and optic fiber placement.

Immunofluorescence was used to confirm expression of GRAB_DA_ in the NAc. Floating coronal sections were washed 3 times in 1x PBS for 30 min and then blocked for 1 – 1.5 hr at room temperature in a solution of 3% normal goat serum (Thermofisher Scientific) and 0.3% Triton X-100 (Thermofisher Scientific) dissolved in PBS. Sections were then washed 3 times in PBS for 15 min and incubated in blocking solution containing chicken anti-GFP polyclonal antibody (1:1000; Abcam, Cambridge, MA) with gentle agitation at 4°C for 18 – 22 hr. Sections were next rinsed 3 times in PBS for 30 min and incubated with goat anti-chicken IgY, Alexa Fluor 488 conjugate (1:500; Abcam) in blocking solution at room temperature for 2 hr. Sections were washed a final 2 times in PBS for 10 min.

tdTomato fluorescence with a Th costain was used to confirm expression of ArchT-tdTomato in VTA_DA_ neurons. Floating coronal sections were washed 3 times in 1x PBS for 30 min and then blocked for 2 hr at room temperature in a solution of 3% normal donkey serum (Thermo Fisher Scientific) and 0.2% Triton X-100 dissolved in PBS. Sections were then washed 3 times in PBS for 15 min and incubated in blocking solution containing rabbit anti-TH antibody (1:1000; EMD Millipore, Burlington, MA) with gentle agitation at 4°C for 44 - 48 hr. Sections were next rinsed 3 times in PBS for 30 min and incubated with goat anti-rabbit IgG, Alexa Fluor 488 conjugate (1:500; Thermofisher Scientific) in blocking solution at room temperature for 2 hr. Sections were washed a final 2 times in PBS for 10 min. Immunofluorescence was also used to confirm expression of ArchT-tdTomato in axons and terminals in the NAc. Floating coronal sections were washed 2 times in 1x PBS for 10 min and then blocked for 2 hr at room temperature in a solution of 10% normal goat serum and 0.5% Triton X-100 dissolved in PBS. Sections were then washed 3 times in PBS for 15 min and incubated in blocking solution containing rabbit anti DsRed polyclonal antibody (1:1000; EMD Millipore, Burlington, MA) with gentle agitation at 4°C for 18 - 22 hr. Sections were next rinsed 3 times in blocking solution for 30 min and incubated with goat anti-rabbit IgG, Alexa Fluor 594 conjugate (1:500; Thermofisher Scientific) in blocking solution at room temperature for 2 hr. Sections were washed a final 2 times in PBS for 10 min.

### Statistical analysis

Datasets were checked for normality using the Kolmogorov-Smirnov test and then analyzed by two-tailed, paired and unpaired Student’s *t* tests, one-, two-, or three-way repeated-measures analysis of variance (ANOVA; GraphPad Prism, GraphPad, San Diego, CA; SPSS, IBM, Chicago, IL). *Post hoc* tests were corrected for multiple comparisons using the Bonferroni method and used to clarify main and interaction effects. To assess the relationship between cue-evoked dopamine and event variables we used multivariate general linear model (Covariance predictor: peak cue-evoked dopamine at CS onset; Dependent variables: test trial, cue-evoked change in lever presses, and cue-evoked change in food-port entries), Pearson and partial correlation (controlling for subject), and linear mixed-model analysis (Repeated factors: CS, trial; Covariance predictor: peak cue-evoked dopamine at CS onset; Random factor: subject; Dependent variable: cue-evoked change in lever presses or food-port entries) (GraphPad Prism, GraphPad, San Diego, CA; SPSS, IBM, Chicago, IL). To evaluate whether cue-evoked dopamine (peak dopamine after CS onset) mediates the influence of CS type on instrumental reward-seeking and reward-checking behavior, we used mediation analysis with the PROCESS package for SPSS, model 4 with 2000 bootstrap samples. We also used this approach to ask whether either reward checking mediates the influence of CS type on reward seeking and vice versa and to ask whether cue-induced reward checking mediates the influence of virus on reward seeking and vice versa. Alpha levels were set at *P* < 0.05.

### Sex as a biological variable

Male and female rats were used in approximately equal numbers for each cohort of each experiment. Individual data points are visually disaggregated by sex. We did not detect any significant effect of sex or interaction between sex and variables of interest (lowest *P* = 0.09).

### Rigor and reproducibility

Group sizes were estimated *a priori* based on prior work using male Long Evans rats in this behavioral task (Malvaez et al., 2015; Lichtenberg and Wassum, 2016; Lichtenberg et al., 2017; Sias et al., 2021; Sias et al., 2024) and to ensure counterbalancing of cue-reward probability pairings. Investigators were not blinded to viral group because they were required to administer virus. All behaviors were scored using automated software (MedPC). Each experiment included at least 2 replication cohorts and cohorts were balanced by viral group, cue-reward probability pairings, sex etc. prior to the start of the experiment.

### Data and code availability

All data that support the findings of this study are available as source data provided with this publication and from the corresponding author upon request. Custom-written MATLAB code is accessible via Dryad repository (https://zenodo.org/records/4926719) (Sias et al., 2021) and available from the corresponding author upon request.

## RESULTS

### Reward prediction shapes reward-pursuit strategy

We first monitored NAc dopamine release as cues influenced reward-pursuit behaviors (Figure 1a-c). Male and female rats were food restricted and received Pavlovian conditioning. We used short-delay conditioning to encourage imminent reward prediction and conflicting instrumental reward-seeking and reward-checking strategies at test (Ostlund and Marshall, 2021). To evaluate the influence of reward prediction on both behavior and dopamine, we used two distinct, 10-s auditory cues (conditioned stimuli, CS) each of which predicted food-pellet reward with a different probability. The high-probability cue (CS_90_) co-terminated with reward delivery on 90% of the trials. The low-probability cue (CS_30_) co-terminated with reward on only 30% of the trials. Rats learned these differential reward-probability predictions. By the end of training, they showed more conditional checks of (i.e., entries into) the food-delivery port during the high-probability cue than the low-probability cue (Figure 1d). To provide rats an instrumental reward-seeking action, they were next trained to lever press to earn food-pellet rewards. All rats acquired the instrumental behavior (final average press rate = 17.27, s.e.m. = 2.39). To ask how the cues affected instrumental seeking v. checking reward-pursuit strategies, we gave rats a Pavlovian-to-instrumental transfer (PIT) test. At test, each cue was presented in pseudorandom order to assess its influence over lever pressing and checks of the food-delivery port. No rewards were delivered, such that cued reward predictions decreased across the test. We operationalized cue-induced instrumental reward seeking as lever presses during the cue. We defined reward checking as Pavlovian conditional entries into the food-delivery port that occurred independent of an instrumental press-enter sequence, thereby excluding food-port entries directly linked to lever presses (see Extended Data Figure 1-2) (Marshall et al., 2023).

**Figure 1.**
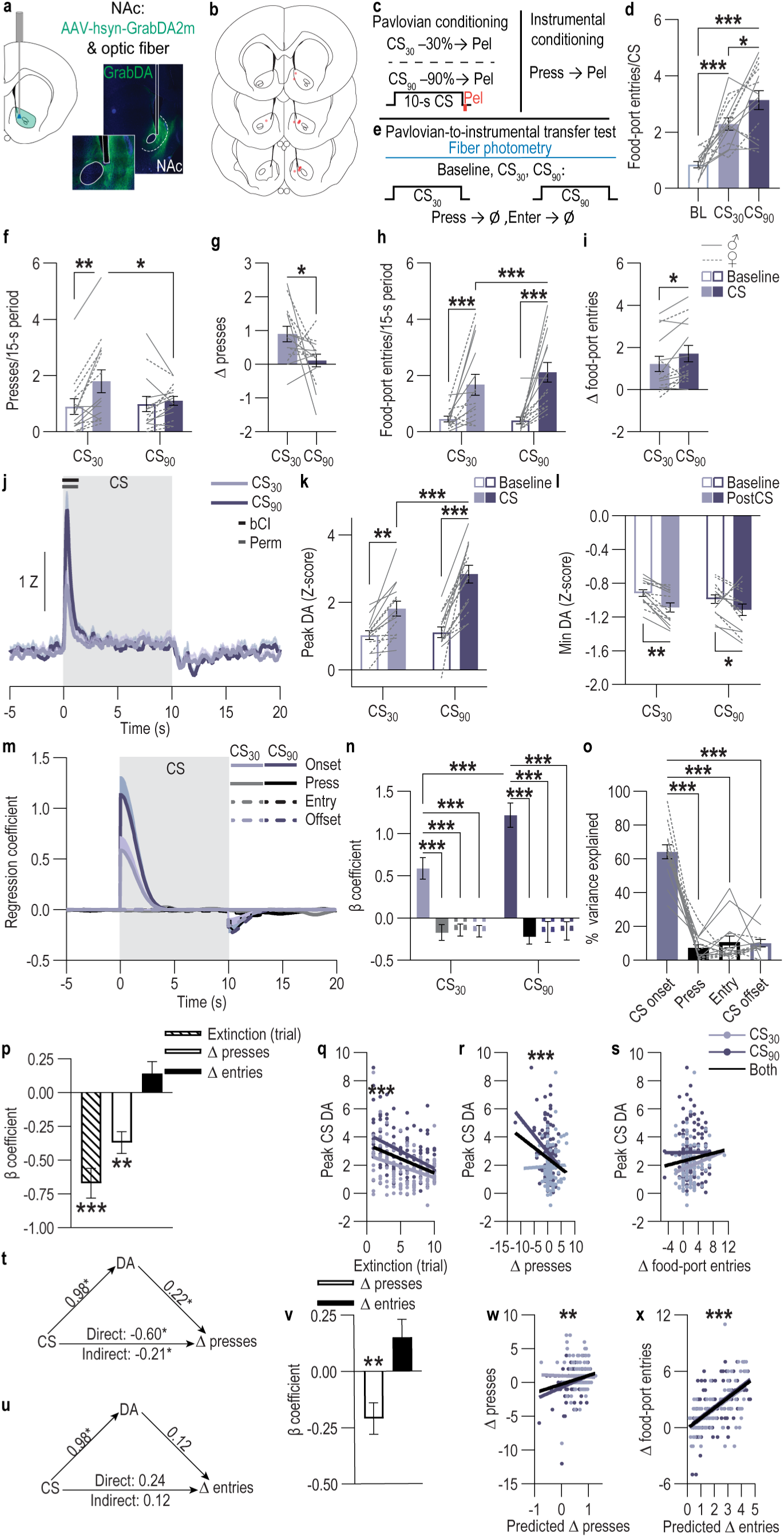
Cue-evoked NAc dopamine is positively modulated by reward prediction and inversely relates to instrumental reward seeking. ***A,*** Left, fiber photometry approach for imaging GRAB_DA_ fluorescence changes in NAc core neurons. Right, Representative immunofluorescence image of NAc GRAB_DA2m_ expression and fiber placement. ***B,*** Fiber placement map. ***C,*** Pavlovian and instrumental conditioning schematic. CS, 10-s conditioned stimulus (Click or Tone) coterminating with food-pellet (Pel) reward on 30% (CS_30_) or 90% (CS_90_) of the trials. Press, lever press earns food pellet (Pel) on a random-interval reinforcement schedule. ***D,*** Food-port entries per 10-s baseline (BL) and CS period averaged across the last 2 days of Pavlovian conditioning. 1-way repeated measures (RM)-ANOVA, F_2,26_ = 24.18, *P* < 0.0001. ***E,*** Pavlovian-to-instrumental transfer test schematic. Ø, no rewards were delivered at test. ***F,*** Lever presses for the 15-s CS (10-s CS + 5-s postCS) and preceding 15-s baseline periods. 2-way RM-ANOVA, CS presence x CS type: F_1,13_ = 4.83, *P* = 0.047; CS presence: F_1,13_ = 21.48, *P* = 0.0005; CS type: F_1,13_ = 2.50, *P* = 0.14. ***G,*** Cue-evoked change (Δ) in lever presses from baseline. 2-tailed t-test, t_13_ = 2.20, *P* = 0.047, 95% Confidence interval (CI) −1.56 to −0.01. ***H,*** Food-port entries, excluding those directly linked to lever presses, for the 15-s CS and baseline periods. 2-way RM-ANOVA, CS presence x CS type: F_1,13_ = 7.69, *P* = 0.02; CS presence: F_1,13_ = 16.40, *P* = 0.001; CS type: F_1,13_ = 2.56, *P* = 0.13. ***I,*** Cue-evoked change in food-port entries. 2-tailed t-test, t_13_ = 2.77, *P* = 0.02, 95% CI 0.11 to 0.88. ***J,*** Cue-evoked GRAB_DA_ fluorescence changes (Z-score). Waveform analysis: bCI, time points where the adjusted 95% bootstrap confidence interval for the difference between CS_30_ and CS_90_ excludes zero (10-sample cluster minimum); Perm, time points for which CS_30_ is significantly (*P* < 0.05) different from CS_90_ from permutation test. ***K,*** Peak CS-onset-evoked GRAB_DA_ (DA) Z-score compared to preCS baseline. 2-way RM-ANOVA, CS onset x CS type: F_1,13_ = 13.88, *P* = 0.003; CS onset: F_1,13_ = 49.91, *P* < 0.0001; CS type: F_1,13_ = 23.25, *P* = 0.0003. ***L,*** Minimum (min) CS offset GRAB_DA_ Z-score relative to baseline before CS offset. 2-way RM-ANOVA, CS offset: F_1,13_ = 20.20, *P* = 0.0006; CS type: F_1,13_ = 1.08, *P* = 0.32; CS offset x CS type: F_1,13_ = 0.56, *P* = 0.47. ***M-O,*** Linear regression model using convolved CS onset, offset, lever presses, and food-port entries to predict GRAB_DA_ signal at each time point 5 s prior to and 20 s after each cue onset from the event kernels. Average regression coefficient of each kernel around cue presentation (m), β coefficients from model (n; 2-way RM-ANOVA, CS type x Event type: F_3,39_ = 5.57, *P* = 0.003; Event type: F_3,39_ = 49.25, *P* < 0.0001; CS type: F_1, 13_ = 3.32, *P* = 0.09), and percentage of variance explained by each event kernel (o; 1-way RM-ANOVA, F_3, 39_ = 62.06, *P* < 0.0001). ***P,*** β coefficients from multivariate general linear model with covariance predictor: peak cue-onset-evoked dopamine, dependent variables: test trial, cue-evoked change in lever presses, and cue-evoked change in food-port entries (excluding those directly linked to presses). Overall model: F_3,254_ = 15.79, *P* < 0.001, η² = 0.16; Extinction test trial: F_3,129_ = 40.05, *P* < 0.001, η² = 0.14; β = −0.67, SE = 0.11, t_256_= −6.33, *P* < 0.001, 95% CI −0.88 - −0.46; Presses: F_1,129_ = 10.88, *P* = 0.001, η² = 0.04; β = - 0.37, SE = 0.08, t_256_= −3.30, *P* = 0.001, 95% CI −0.44 - −0.11; Food-port entries: F_1,129_ 2.54, *P* = 0.11, η² = 0.01; β = 0.14, SE = 0.09, t_256_= 1.59, *P* = 0.11, 95% CI −0.03 - 0.32. ***Q-S,*** Relationship between peak Cue-evoked GRAB_DA_ Z-score and test trial (q; 2-tailed Partial correlation, controlling for subject, r_261_ = −0.37, *P* < 0.001.), cue-evoked change in lever presses (r; 2-tailed Partial correlation, r_255_ = −0.20, *P* < 0.001), or food-port entries (s; 2-tailed Partial correlation, r_255_ = −0.12, *P* = 0.06). ***T,*** Mediation analysis. CS (CS_30_→CS_90_) effect on cue-onset-evoked dopamine: β = 0.98, SE = 0.19, t_256_= 5.15, *P* < 0.001, 95% CI 0.61 – 1.36; effect of Cue-evoked dopamine on Cue-evoked presses: β = −0.22, SE = 0.09, t_256_= −2.49, *P* = 0.01, 95% CI −0.38 – −0.05; Direct effect of CS on presses: β = −0.60, SE = 0.28, t_256_= −2.18, *P* = 0.03, 95% CI −1.15 – −0.06; Indirect effect of CS on presses mediated by dopamine: β = −0.21, BootSE = 0.13, BootCI - 0.51 – −0.06. ***U,*** Mediation analysis. CS (CS_30_→CS_90_) effect on cue-evoked dopamine: β = 0.98, SE = 0.19, t_256_= 5.15, *P* < 0.001, 95% CI 0.61 – 1.36; effect of cue-evoked dopamine on Cue-evoked food-port entries (excluding those directly linked to lever pressing): β = 0.11, SE = 0.09, t_256_= 1.27, *P* = 0.20, 95% CI −0.06 – 0.30; Direct effect of CS on entries: β = 0.23, SE = 0.29, t_256_= 0.80, *P* = 0.43, 95% CI - 0.35 – 0.83; Indirect effect of CS on presses mediated by dopamine: β = 0.12, BootSE = 0.09, BootCI −0.04 – 0.32. ***V,*** β coefficients from linear mixed models with covariance predictor: peak Cue-evoked dopamine, dependent variable: cue-evoked change in lever presses, repeated factors: CS, trial, and random factor: subject (β = −0.21, standard error (SE) = 0.07, t_136.60_ = −3.06, *P* = 0.003, 95% CI −0.35 - - 0.08 and covariance predictor: peak Cue-evoked dopamine, dependent variable: cue-evoked change in food-port entries, repeated factors: CS, trial, and random factor: subject (β = 0.15, SE = 0.08, t_139.73_ = 1.89, *P* = 0.06, 95% CI −0.007 - 0.30). ***W-X,*** Correlation between actual cue-evoked change in presses (v; Pearson r_258_ = 0.20, *P* = 0.001) or food-port entries (w; Pearson r_258_ = 0.59, *P* < 0.0001) and that predicted by cue-evoked dopamine using the linear mixed model. Data presented as mean ± s.e.m. Males = solid lines, Females = dashed lines. \**P* < 0.05, \*\**P* < 0.01, \*\*\**P* < 0.001, Bonferroni corrected.

Whereas cues that predict reward with low probability bias towards instrumental reward seeking, highly predictive cues bias towards checking for the expected reward (Marshall et al., 2020). Indeed, at test, the low-probability cue increased lever pressing (Figure 1f-g). Such instrumental reward seeking was not triggered in response to the high-probability cue. Instead, this cue preferentially elicited a reward-checking strategy. The high-probability cue caused more conditional entries into the food-delivery port than the low-probability cue (Figure 1h-i). Because rats cannot readily physically lever press and check the food port at the same time, we conducted 4 analyses to assess whether overt response competition explains the differential performance of instrumental reward seeking v. reward checking during the low- v. high-probability cue. First, if response competition explains differential instrumental reward-seeking v. reward-checking behavior, then the cue-induced changes in lever presses and food-port entries should be significantly inversely correlated. They were not (Partial correlation, controlling for subject, r_255_ = −0.08, *P* = 0.19; within subject across trials Pearson correlation, r_20_ = −0.39, *P* = 0.09; between subjects Pearson correlation, r_28_ = −0.10, *P* = 0.60; Partial correlation, controlling for subject using only the 10-s CS periods, r_255_ = −0.04, *P* = 0.55). Second, if response competition explains the results, performance of one behavior should decrease the probability of another. It did not. During the cues, a food-port entry did not decrease the probability of a lever press, nor did a lever press decrease food-port-entry probability (Extended Data Figure 1-3a-b). Third, if response competition explains the differential instrumental reward-seeking v. reward-checking during the low- v. high-probability cue, then a high reward-checking response should preclude the low-probability cue from also promoting instrumental reward seeking. It did not. The low-probability cue caused greater instrumental reward seeking than the high-probability cue even on trials for which the cues triggered high (≥ 1 standard deviation above the mean) reward-checking behavior (Extended Data Figure 1-3c). Lastly, either did the cue-induced change in reward checking significantly mediate the influence of the CS type on instrumental reward seeking nor did the cue-induced change in reward seeking significantly mediate the influence of CS type on reward checking (Extended Data Figure 1-3d-e). Thus, similar to prior reports (Marshall et al., 2023), although instrumental reward seeking and reward checking are competing responses, their differential performance during the low- v. high-probability cues was not due to overt response competition. There were modest changes in elicited behavioral strategy across test trials (Extended Data Figure 1-4a-b). Most notably, initially, the high-probability cue suppressed instrumental reward seeking, but this seeking response increased as reward prediction weakened across trials. Thus, reward prediction shapes reward-pursuit strategy. Cues that signal imminent reward with high probability preferentially elicit a strategy of checking for the expected reward. Conversely, cues that signal reward with low probability bias towards instrumental reward seeking.

### Cue-evoked NAc dopamine is positively modulated by reward prediction and inversely relates to instrumental reward seeking

We exploited this task to ask the extent to which cue-evoked NAc dopamine release is shaped by reward prediction and relates to reward-pursuit behaviors. We used fiber photometry to record fluorescent activity of the G-protein-coupled-receptor-activation-based dopamine sensor (GRAB_DA_) in the NAc core during the PIT test (Fig. 1a-c,e). The onset of both cues triggered robust, transient dopamine release (Figure 1j-k, see Extended Data Figure 1-5 for non-subtracted GRAB_DA_ fluorescence response). We also detected a dopamine dip at cue offset, when the predicted reward would have been delivered, but was omitted (Figure 1j,l), and transient dopamine elevations prior to instrumental reward-seeking and checking behaviors during the cues (Extended Data Figure 1-6). If dopamine is shaped by reward prediction, the high-probability cue should trigger more dopamine release than the low-probability cue. Indeed, the cue-evoked dopamine response was larger for the high- than low-probability cue (Figure 1j-k). If cue-evoked dopamine is shaped by reward prediction, then it should also have declined as the reward prediction weakened across trials of the unrewarded test. This was the case (Figure 1n; Extended Data Figure 1-4c). Because lever pressing and food-port entries can freely occur during cue presentation, we used a linear regression model to parse the extent to which dopamine fluctuations are explained by the cues themselves v. concomitant behavioral responses (Parker et al., 2016; Greenstreet et al., 2025). The model included cue onset and offset, lever presses, and food-port entries. Dopamine release during the cues was best explained by the CS onset kernel, indicating that the differential dopamine response to the high- v. low-probability cue was due to the cues themselves rather than the differential behavioral responses they triggered (Figure 1m-n). The influence of CS on dopamine was stronger for the high-probability cue than the low-probability cue. Thus, cue-evoked NAc dopamine is shaped by reward prediction occurring at cue onset. Stronger reward predictions cause greater cue-evoked NAc dopamine release.

We next further probed the relationship between cue-evoked NAc dopamine and reward prediction and asked how this signal relates to the influence of cues on reward-pursuit strategy. We first used a multivariate general linear model including data from each trial for each subject to ask the extent to which cue-evoked dopamine, measured by the peak dopamine response to cue onset, relates to the declining reward prediction resulting from extinction across the test (defined as test trial) and cue-induced changes in instrumental reward-seeking and reward-checking behavior (Figure 1p). Cue-evoked dopamine significantly related to this multivariate combination. If dopamine generally promotes reward pursuit, it should positively relate to both instrumental reward-seeking and reward-checking behaviors, or perhaps just instrumental reward seeking (du Hoffmann and Nicola, 2014; Hamid et al., 2015; Syed et al., 2016). If, however, dopamine supports reward prediction to shape reward-pursuit strategy, it should inversely relate to test trial and differentially relate to instrumental reward-seeking v. reward-checking behavior. The data support the latter. Cue-evoked dopamine significantly inversely related to test trial (Figure 1p,q), indicating that as reward prediction weakened, so too did the cue-evoked dopamine. Thus, cue-evoked dopamine tracks trial-dependent changes in reward prediction. Cue-evoked dopamine also significantly inversely related to cue-induced instrumental reward seeking (Figure 1p,r), but did not significantly relate to reward-checking behavior (Figure 1p,s). The negative relationship between cue-evoked dopamine and cue-evoked instrumental reward seeking held true both between- and within-subjects (Extended Data Figure 1-7). Subjects for which the cues elicited a larger dopamine response showed less cue-evoked instrumental reward seeking. On trials in which the cue elicited a larger dopamine response it also tended to trigger less instrumental reward seeking. Thus, larger cue-evoked dopamine is associated with stronger reward predictions and less instrumental reward seeking. We next used mediation analyses to ask whether cue-evoked dopamine mediates the influence the cued reward prediction on reward-pursuit strategy. We found that the magnitude of the peak cue-evoked dopamine response explains a meaningful portion of the influence the cued reward prediction (CS_30_ v. CS_90_) on instrumental reward seeking (Figure 1t). Dopamine did not significantly mediate the influence of CS type on reward checking (Figure 1u). Lastly, we asked whether the peak cue-evoked dopamine response could predict cue-induced behavior. Using a linear mixed-model analysis, we were able to reliably inversely predict instrumental reward seeking from cue-evoked dopamine (Figure 1w,v; Extended Data Figure 1-4d), indicating larger cue-evoked NAc dopamine predicts less instrumental reward seeking. Cue-evoked reward checking was positively, though not statistically significantly, predicted from cue-evoked dopamine (Figure 1v,x; Extended Data Figure 1-4e). Together, these results indicate that cue-evoked NAc dopamine is positively modulated by reward prediction and inversely relates to instrumental reward seeking. Stronger cue-evoked reward predictions elicit more NAc dopamine release and this is associated with less instrumental reward seeking.

### Cue-evoked NAc core dopamine mediates the influence of cued reward prediction on reward pursuit

We next asked whether cue-evoked NAc dopamine release mediates the influence of cued reward prediction on reward-pursuit behaviors. Because we found cue-evoked dopamine is positively modulated by reward prediction and inversely relates to instrumental reward seeking, we reasoned that cue-evoked NAc core dopamine may support reward prediction to shape reward-pursuit strategy. Large cue-evoked NAc core dopamine may support high imminent reward prediction to constrain instrumental reward seeking in favor of checking for the strongly predicted reward. This leads to the intriguing, perhaps surprising, prediction that inhibition of cue-evoked NAc dopamine release should increase instrumental reward seeking and decrease reward checking.

We tested this using optogenetic inhibition of cue-evoked NAc dopamine release during a PIT test (Figure 2a-c). We cre-dependently expressed the inhibitory opsin archaerhodopsin T (ArchT) bilaterally in ventral tegmental area dopamine (VTA_DA_) neurons of male and female tyrosine hydroxylase (Th)-cre rats (Figure 2a-b) and implanted optical fibers bilaterally over NAc core (Figure 2b). For controls, we either cre-dependently expressed tdTomato bilaterally in VTA_DA_ neurons of Th-cre rats or infused a control mCherry AAV in wildtype (Th-cre-) rats and, in both cases, implanted optical fibers bilaterally over NAc core. Rats were food restricted and received Pavlovian short-delay conditioning, as above, during which two distinct, 10-s auditory cues each predicted a food-pellet reward with either high (90%) or low (30%) probability. Rats learned these differential reward-probability predictions (Extended Data Figure 2-1a). Rats then received instrumental conditioning to lever press to earn food-pellet rewards (Extended Data Figure 2-1b). They were then given a PIT test during which the lever was available and each cue was presented in pseudorandom order to assess its influence over instrumental reward-seeking lever presses and checks of the food-delivery port. No rewards were delivered at test. We optically inhibited (532 nm, 10 mW, 2.5 s) VTA_DA_ axons and terminals in the NAc coincident with each cue during this test. We restricted optical inhibition to cue onset to inhibit only the phasic cue-induced dopamine elevation. Reward prediction influenced reward-pursuit strategy in controls. Only the low-probability cue increased instrumental reward seeking (Figure 2d-e). The high-probability cue increased reward checking more than the low-probability cue (Figure 2f-g). Optical inhibition of cue-evoked VTA_DA_→NAc activity elevated the instrumental reward-seeking response to both cues (Figure 2d-e) and decreased cue-evoked reward checking during the high-probability cue (Figure 2f-g; see Extended Data Figure 2-2 for data on all entries and press-linked entries). These effects were not correlated (two-tailed, between-subject Pearson r, Control, CS_30_: r_12_ = 0.27, *P* = 0.39; CS_90_: r_12_ = −0.10, *P* = 0.76; Both CSs: CS_30_: r_24_ = −0.01, *P* = 0.95; ArchT, CS_30_: r_14_ = 0.22, *P* = 0.44; CS_90_: r_14_ = −0.18, *P* = 0.55; Both CSs: CS_30_: r_28_ = −0.03, *P* = 0.88). Moreover, the cue-induced change in reward checking did not significantly mediate the influence of Group (control→ArchT) on cue-induced instrumental reward seeking (Indirect effect of Group on cue-induced food-port entries mediated by cue-induced presses: β = 0.01, BootSE = 0.09, BootCI −0.15 – 0.22) and neither did cue-induced reward seeking significantly mediate the influence of the Group on reward checking (Indirect effect of Group on cue-induced food-port entries mediated by cue-induced presses: β = −0.009, BootSE = 0.06, BootCI −0.17 – 0.11). Thus, the VTA_DA_→NAc inhibition-induced increase in instrumental reward seeking was not a secondary consequence of the decrease in reward checking or vice versa. Despite inhibition occurring only at cue onset, these effects persisted throughout the duration of the cue period (Extended Data Figure 2-3). Thus, cue-evoked, transient, NAc core dopamine release mediates the influence of reward prediction on reward-pursuit strategy, which, in this case of imminent reward prediction, opposes instrumental reward seeking and promotes checking for a highly predicted reward.

**Figure 2.**
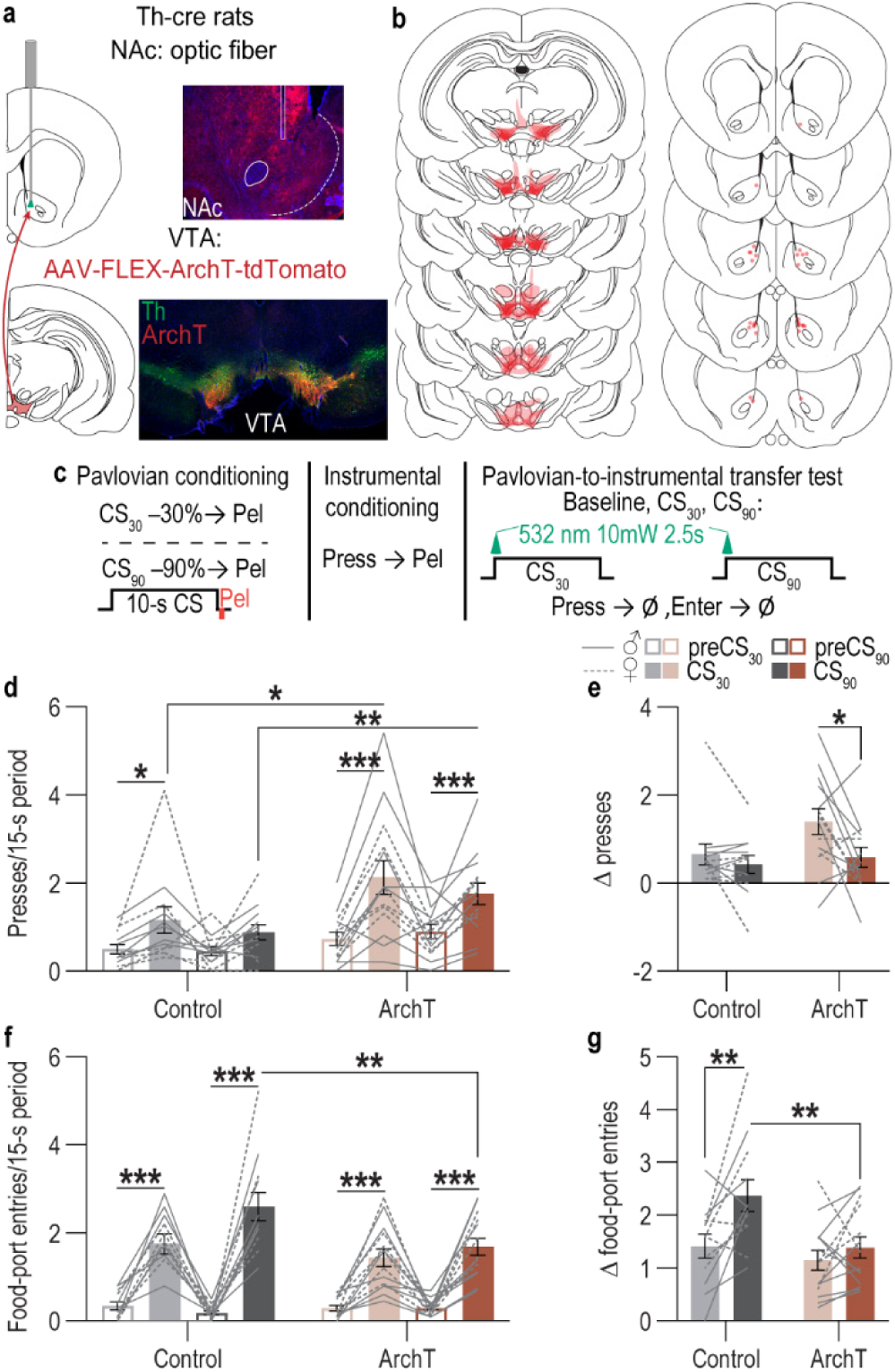
Cue-evoked NAc core dopamine constrains instrumental reward seeking and promotes reward checking. ***A,*** Left, optogenetic approach for bilateral inhibition of VTA_DA_→NAc projections. Right top, representative image of fiber placement in the vicinity of immunofluorescent ArchT-tdTomato-expressing VTA_DA_ axons and terminals in NAc. Right bottom, representative immunofluorescent image of ArchT-tdTomato expression in VTA_DA_ (Th+) neurons. ***B,*** VTA ArchT expression and NAc fiber placement maps. ***C,*** Task training and test schematic. CS, 10-s conditioned stimulus (Click or Tone) coterminating with food pellet (Pel) reward on 30% (CS_30_) or 90% (CS_90_) of the trials. Press, lever press earns food-pellet (Pel) reward on a random-interval reinforcement schedule. Ø, no rewards were delivered. ***D,*** Lever presses for the 15-s CS (10-s CS + 5-s postCS) and preceding baseline periods. 3-way RM-ANOVA, Virus x CS presence: F_1,24_ = 4.48, *P* = 0.04; Virus: F_1,24_ = 6.75, *P* = 0.02; CS presence: F_1,24_ = 36.92, *P* < 0.0001; CS presence x CS type: F_1,24_ = 3.44, *P* = 0.07; CS type: F_1,24_ = 1.07, *P* = 0.31; Virus x CS type: F_1,24_ = 0.06, *P* = 0.81; Virus x CS presence x CS type: F_1,24_ = 0.57, *P* = 0.46. ***E,*** Cue-evoked change (Δ) in lever presses from baseline. 2-way RM-ANOVA, CS type: F_1,24_ = 6.33, *P* = 0.02; Virus: F_1,24_ = 2.58, *P* = 0.12; Virus x CS type: F_1,24_ = 1.97, *P* = 0.17. ***F,*** Food-port entries, excluding those directly linked to lever presses, for the 15-s CS and preceding baseline periods. 3-way RM-ANOVA, Virus x CS presence: F_1,24_ = 5.68, *P* = 0.03; CS presence x CS type: F_1,24_ = 9.90, *P* = 0.004; Virus x CS presence x CS type: F_1,24_ = 3.72, *P* = 0.07; CS presence: F_1,24_ = 137.90, *P* < 0.0001; CS type: F_1,24_ = 5.86, *P* = 0.02; Virus: F_1,24_ = 3.82, *P* = 0.06; Virus x CS type: F_1,24_ = 1.22, *P* = 0.28. ***G,*** Cue-evoked change in food-port entries. 2-way RM-ANOVA, Virus: F_1,24_ = 5.66, *P* = 0.03; CS type: F_1,24_ = 10.18, *P* = 0.004; Virus x CS type: F_1,24_ = 3.65, *P* = 0.07. Control, *N* = 12 rats (6 male; 9 Th-cre+, 3 TH-cre-) ArchT, *N* = 14 Th-cre+ rats (8 male). Data presented as mean ± s.e.m. Males = solid lines, Females = dashed lines. \**P* < 0.05, \*\**P* < 0.01, \*\*\**P* < 0.001, Bonferroni corrected.

## DISCUSSION

Here we investigated whether cue-evoked dopamine supports reward prediction to shape reward-pursuit strategy. We found that cue-evoked NAc core dopamine is positively shaped by reward prediction and inversely relates to and predicts instrumental reward seeking. Cues that predict imminent reward with high probability trigger a large dopamine response and Pavlovian checking for the expected reward in the delivery location, rather than instrumental reward seeking. Cues that predict reward with low probability elicit less dopamine and bias towards instrumental seeking, rather than checking for reward. Correspondingly, NAc dopamine mediates the influence of reward prediction on reward-pursuit strategy. Cue-evoked NAc dopamine release to short-delay cues opposes instrumental reward seeking and promotes checking for a highly predicted reward. Thus, cue-evoked NAc dopamine supports high imminent reward prediction to constrain instrumental reward seeking in favor of a Pavlovian reward-checking strategy.

Reward prediction shapes how reward is pursued. Consistent with prior reports (Marshall et al., 2020), we found that cues that signal imminent reward with high probability tend to elicit a strategy of checking for the expected reward in its delivery location. Conversely, low-probability cues preferentially trigger instrumental reward seeking. Differential performance of instrumental reward-seeking v. reward-checking forms of reward pursuit was not simply due to overt response competition (Marshall et al., 2023). Rather they reflect distinct, yet conflicting, global v. focal reward-pursuit strategies (Timberlake, 1994; Marshall et al., 2023). Reward-pursuit strategies were selected on-the-fly in a novel situation, without than prior reinforcement, indicating they were shaped by reward predictions. Cue-evoked reward predictions may, therefore help to flexibly resolve conflict between these competing response tendencies (Marshall et al., 2023). Our results are consistent with evidence that cue-evoked predictions of the probability (Marshall et al., 2020), timing (Lovibond, 1981; Crombag et al., 2008; Marshall and Ostlund, 2018), and value (Marshall et al., 2023) of reward regulate reward-pursuit strategy.

That strong reward prediction constrains instrumental reward seeking in favor of checking for the expected reward is particularly the case for cues that signal imminent, non-contingent reward (Konorski, 1967; Ostlund and Marshall, 2021), as we used here. Indeed, long-duration cues that set the occasion for sporadic, infrequent reward tend to promote instrumental reward seeking (Crombag et al., 2008; Wassum et al., 2013; Collins et al., 2016; Halbout et al., 2019). The one-outcome/one-lever design here limits our ability to understand whether reward pursuit was driven by the predicted reward identity (i.e., specific PIT), more general motivational properties (i.e., general PIT), or, perhaps, a process unique to cues that signal imminent reward (Ostlund and Marshall, 2021; Marshall et al., 2023). Towards the latter, whereas both specific and general PIT to loosely predictive cues are insensitive to devaluation of the predicted reward (Colwill and Rescorla, 1990; Rescorla, 1994; Holland, 2004), reward devaluation causes high-probability, imminent reward-predicting cues that would normally suppress instrumental reward seeking to promote such behavior (Marshall et al., 2023). Thus, the psychological processes through which short-delay cues influence reward pursuit and the extent to which they differ from long-delay, loosely-predictive cues are important questions for future investigation. Such studies would also benefit from asking whether the balance between instrumental reward seeking and reward checking is shaped not only by cue-reward predictions but also the strength and nature of the instrumental association.

NAc core dopamine is positively modulated by cue-evoked reward prediction. Consistent with prior reports (Roitman et al., 2004; Day et al., 2007; Day et al., 2010; Flagel et al., 2011; Wassum et al., 2013; Aitken et al., 2015; Saddoris et al., 2015; Collins et al., 2016; Sun et al., 2018; Kutlu et al., 2021; Garr et al., 2024; Bornhoft et al., 2025), we found that reward-predictive cues elicit phasic dopamine release in the NAc core. We discovered that this signal is positively shaped by reward prediction. Cues that signal reward with high probability elicit more NAc dopamine than low-probability cues and cue-evoked dopamine tracks trial-dependent changes in reward prediction. This accords with prior evidence of reward probability encoding in both VTA dopamine neuron firing (Fiorillo et al., 2003; Tobler et al., 2005; Eshel et al., 2016) and NAc dopamine release (Hart et al., 2015; Garr et al., 2024) and with evidence that cue-evoked NAc dopamine is modulated by predicted reward value (Gan et al., 2009; Day et al., 2010; Aitken et al., 2016; Syed et al., 2016). An important future question is the extent to which dopamine is modulated by other features of the predicted reward. Whereas probability and general value shape NAc cue-evoked dopamine, specific features of the predicted reward may not (Halbout et al., 2019; Taira et al., 2024), at least in NAc core (Sias et al., 2024). Whether dopamine’s modulation by reward prediction is associated with its signaling of prediction error (Schultz et al., 1997; Steinberg et al., 2013; Hart et al., 2014) and/or salience (Kutlu et al., 2021; Kutlu et al., 2022), is also ripe for investigation.

Cue-evoked NAc dopamine mediates the influence of reward prediction on reward-pursuit strategy. We found that cue-evoked NAc core dopamine was not only shaped by reward prediction, but also inversely related to and predicted the ability of the cue to promote instrumental reward seeking. Larger cue-evoked dopamine is associated with stronger reward predictions and less instrumental reward seeking. Correspondingly, we found that inhibition of cue-evoked NAc dopamine increased instrumental reward seeking. It also decreased reward checking, though it did not fully suppress this behavior, consistent with prior evidence (Lex and Hauber, 2008; Flagel et al., 2011; Saunders and Robinson, 2012; Halbout et al., 2019). Therefore, the dopamine release triggered in the NAc core by short-delay cues constrains instrumental reward seeking and promotes the reward-checking strategy. Despite only brief inhibition at cue onset, the behavioral effects persisted during and immediately after the cue. Thus, the cue-evoked dopamine transients can support a transition in the reward-prediction state (Kalmbach et al., 2022) that controls subsequent reward-pursuit strategy, in this case constraining instrumental reward seeking and promoting conditional reward checking. Given that NAc dopamine is shaped by reward prediction, its inhibition may disrupt reward prediction, thereby causing selection of the reward-pursuit strategy more consistent with a low probability of non-contingent reward: instrumental reward seeking. It is also plausible that this manipulation removed the motivational value of the reward prediction leaving other reward features, like identity, intact to drive PIT through an associative priming or outcome-response process (Rescorla, 1992; Balleine and Ostlund, 2007; Watson et al., 2018). Indeed, NAc core dopamine is unlikely to be necessary for such identity information (Corbit and Balleine, 2011; Taira et al., 2024). Further arbitration of these possibilities will help refine understanding of the function of NAc core cue-evoked dopamine.

In either case, these results indicate that cue-evoked NAc dopamine mediates the influence of reward prediction on reward-pursuit strategy to flexibly constrain instrumental reward seeking and promote a more focal reward-checking strategy.

Differences between our findings and prior work reveal important possibilities about how NAc dopamine function may differ based on reinforcement history and the duration, strength, and nature of reward prediction. Our evidence that cue-evoked NAc dopamine inversely relates to instrumental reward seeking contrasts with findings that cues elicit more NAc dopamine when they promote incentive motivation (Flagel et al., 2011) or the need to instrumentally seek reward (Syed et al., 2016), than when they encourage waiting at the reward-delivery location. Unlike the present design, in these studies reward-pursuit strategy was reinforced. Reinforcement may cache the value of the action to the predictive cue, such that the cue-evoked dopamine not only supports reward prediction, but also the reinforced motivational value of instrumental reward seeking (Hamid et al., 2015; Syed et al., 2016). Thus, the information dopamine conveys and how it influences reward pursuit may differ if reward-pursuit strategies have been reinforced v. are being determined on-the-fly. Our results also contrast with those indicating that tonic inactivation of NAc dopamine input (Halbout et al., 2019) or receptors (Lex and Hauber, 2008) attenuates instrumental reward seeking in response to long-duration, loosely-predictive reward. This could reflect differences in tonic v. phasic dopamine function (Niv et al., 2007; Schultz, 2007). It could also indicate that dopamine function depends on the duration, strength, and nature of reward prediction. These variables may shape the downstream circuitry engaged by the cue. Whereas, strong, imminent reward prediction may engage circuitry to constrain global, instrumental reward seeking and promote focal reward checking, cues that signal less, delayed, uncertain, and/or infrequent reward may engage distinct circuitry to promote instrumental reward seeking. Cue-evoked dopamine may accentuate whichever reward-pursuit circuit is engaged. Another critical question is whether dopamine functions in regions outside the NAc, like the basolateral amygdala (Sias et al., 2024), dorsal striatum (Ostlund et al., 2011; Bernklau et al., 2024), and medial prefrontal cortex (Floresco et al., 2006), also contribute to reward predictions and shaping reward-pursuit behavior. Dopamine is likely to have unique function based on function of the region in which it is released.

Dopamine, including that elicited by cues (Maes et al., 2020), has long been known to function as a teaching signal to support learning (Waelti et al., 2001; Schultz, 2002; Lerner et al., 2021). Our results add to a growing body of evidence that dopamine does not simply function retrospectively in learning (Schultz, 2002; Glimcher, 2011; Kahnt and Schoenbaum, 2025), but also prospectively to regulate behavior (Howe and Dombeck, 2016; Schultz, 2016; Berke, 2018; Kaźmierczak and Nicola, 2022). They add that NAc core dopamine can support reward prediction to flexibly regulate reward-pursuit strategy. Inability to adaptively regulate reward-pursuit behaviors are symptoms of psychiatric conditions, including depression and addictions, conditions that are also marked by dopamine disruption. Thus, these data also aid our understanding and treatment of motivational symptoms of psychiatric disease.

## AUTHOR CONTRIBUTIONS

KMW, SBO, AS, and MM conceptualized and designed the experiments. MM & KW analyzed and interpreted the data, and wrote the paper. AS executed the fiber photometry experiments and did the initial analysis. NKG and MM conducted the optogenetic experiments and did the initial analysis. KR conducted the linear regression kernel analysis. SBO provided guidance on experimental execution, analysis, and interpretation and contributed to manuscript preparation.

## ACKNOWLEDGEMENTS

This research was supported by NIH R01MH126285 (SBO & KMW), NIH R01DA057084 (KMW), the Staglin Center for Behavior and Brain Sciences, and the Wendell Jeffrey and Bernice Wenzel Term Chair in Behavioral Neuroscience.

## EXTENDED DATA FIGURES

**Extended Data Figure 1-1.**
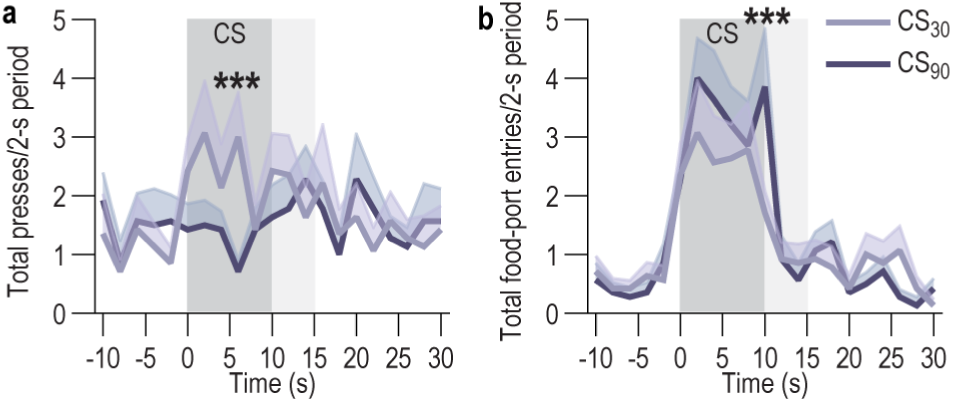
Cues differentially promote instrumental reward seeking v. checking based on reward prediction throughout the cue during Pavlovian-to-instrumental transfer. ***A-B,*** Lever presses *(A)* and food-port entries, excluding those directly linked to lever presses, *(B)* in 2-s bins before, during, and after cue (conditioned stimulus, CS) presentation. Gray, 10-s CS; Light gray, 5-s post-CS period. 2-way RM-ANOVA, Presses, Time x CS type: F_20,260_ = 2.07, *P* = 0.005; Time: F_20,260_ = 1.69, *P* = 0.03; CS type: F_1,13_ = 1.18, *P* = 0.30; Food-port entries, 2-way RM-ANOVA, Time x CS type: F_20,260_ = 2.12, *P* = 0.004; Time: F_20,260_ = 10.98, *P* < 0.0001; CS type: F_1,13_ = 1.41, *P* = 0.26. Data presented as mean ± s.e.m. \*\*\**P* < 0.001, CS_30_ v. CS90, Bonferroni corrected. Cues that signal reward with low probability biased behavior towards instrumental reward seeking. This instrumental reward-seeking strategy was not triggered in response to the high-probability cue. Instead, this cue preferentially elicited a reward-checking strategy (entries into the food-delivery port). Because cue-evoked changes in instrumental reward-seeking and reward-checking behavior persisted into the 5-s post-cue period, when reward was delivered during training, we used a 15-s CS (10-s CS + 5-s postCS) to quantify Cue-evoked behavioral changes during PIT. Reward checking was lower in the post-CS period than the CS period. Instrumental reward seeking was similar between periods. *N* = 14 rats (7 male).

**Extended Data Figure 1-2.**
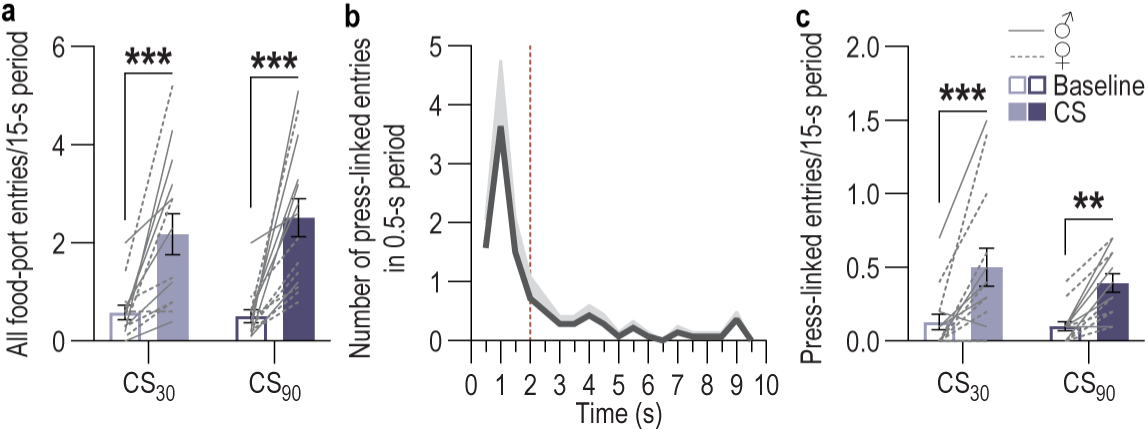
All and press-linked cue-evoked entries into the food-delivery port during Pavlovian-to-instrumental transfer. ***A,*** All food-port entries for the 15-s CS (10-s CS + 5-s postCS) and preceding baseline period. 2-way RM-ANOVA, CS presence x CS type: F_1,13_ = 3.17, *P* = 0.10; CS presence: F_1,13_ = 24.56, *P* = 0.0003; CS type: F_1,13_ = 0.60, *P* = 0.45. Both cues cause conditional entries into the food-delivery port. ***B,*** Distribution of the time of food-port entries occurring directly after a lever press. Many entries into the food-delivery port occurred immediately (within 2-s) directly following a lever press, indicating they are part of an instrumental press-check sequence (Halbout et al., 2019; Halbout et al., 2022). Therefore, in the main figures, to isolate conditional food-port checks from those that occurred as part of an instrumental press-enter sequence, we excluded food-port entries that directly followed a lever press within 2 s. ***C,*** Press-linked food-port entries (entry occurring directly after and within 2-s of a press) for the 15-s CS and preceding baseline periods. 2-way RM-ANOVA, CS presence: F_1,13_ = 34.73, *P* < 0.0001; CS type: F_1,13_ = 0.67, *P* = 0.43; CS presence x CS type: F_1,13_ = 0.62, *P* = 0.44. *N* = 14 rats (7 male). Data presented as mean ± s.e.m. Males = solid lines, Females = dashed lines. \*\**P* < 0.01, \*\*\**P* < 0.001, Bonferroni corrected.

**Extended Data Figure 1-3.**
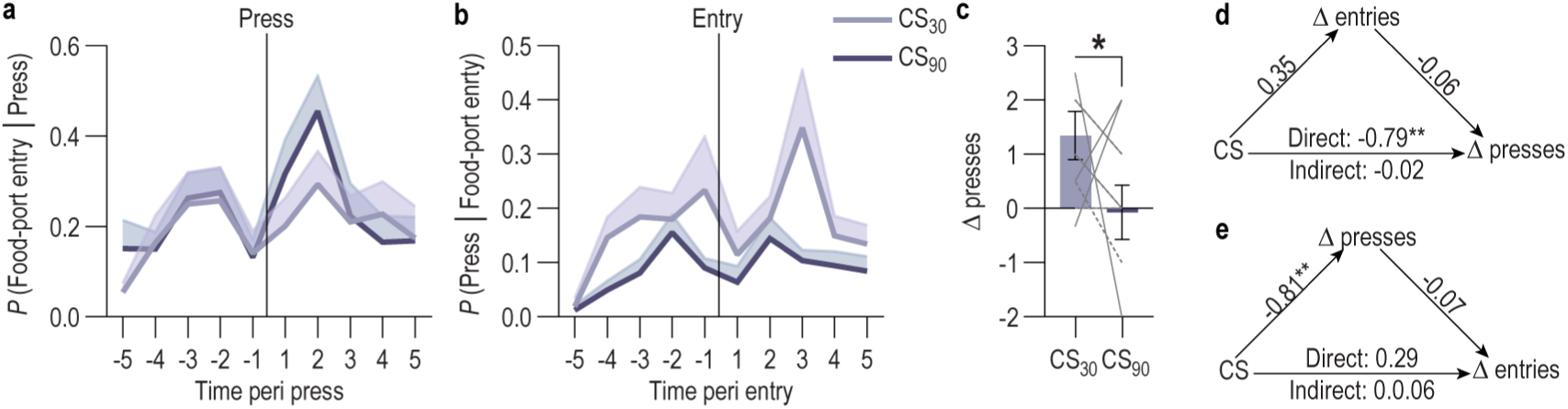
Lever presses do not affect the probability of a food-port entry and vice versa. ***A.*** Probability of food-port entries as a function of time from each lever press during the cues. ***B.*** Probability of lever presses as a function of time from each food-port entry during the cues. ***C.*** Cue-evoked change (Δ) in lever presses from baseline averaged across trials in which entries were ≥ 1 standard deviation higher than the mean for that subject/CS. 2-tailed t-test, t_13_ = 2.60, P = 0.02, 95% Confidence interval (CI) −2.59 to −0.24*. **D,*** Mediation analysis. Cue type (CS_30_→CS_90_) effect on cue-evoked food-port entries (excluding those directly linked to lever presses): β = 0.35, SE = 0.28, t_256_= 1.24, *P* = 0.21, 95% CI −0.21 – 0.91; effect of cue-evoked entries on cue-evoked presses: β = −0.06, SE = 0.06, t_256_= −1.04, *P* = 0.30, 95% CI −0.18 – 0.05; Direct effect of CS type on presses: β = −0.79, SE = 0.26, t_256_= −2.98, *P* = 0.003, 95% CI −1.31– −0.27; Indirect effect of cue type on presses mediated by entries: β = −0.02, BootSE = 0.03, BootCI −0.09 – 0.03. ***E,*** Mediation analysis. cue type (CS_30_→CS_90_) effect on cue-evoked lever presses: β = −0.81, SE = 0.27, t_256_= −3.0, *P* = 0.002, 95% CI −1.34 – −0.29; effect of cue-evoked presses on cue-evoked entries: β = −0.06, SE = 0.07, t_256_= −1.04, *P* = 0.30, 95% CI −0.18 – 0.05; Direct effect of cue type on entries: β = 0.30, SE = 0.29, t_256_= −1.03, *P* = 0.31, 95% CI −0.27 – 0.86; Indirect effect of cue type on entries mediated by presses: β = 0.06, BootSE = 0.06, BootCI −0.04 – 0.19. *N* = 14 rats (7 male). Data presented as mean ± s.e.m. \**P* < 0.05.

**Extended Data Figure 1-4.**
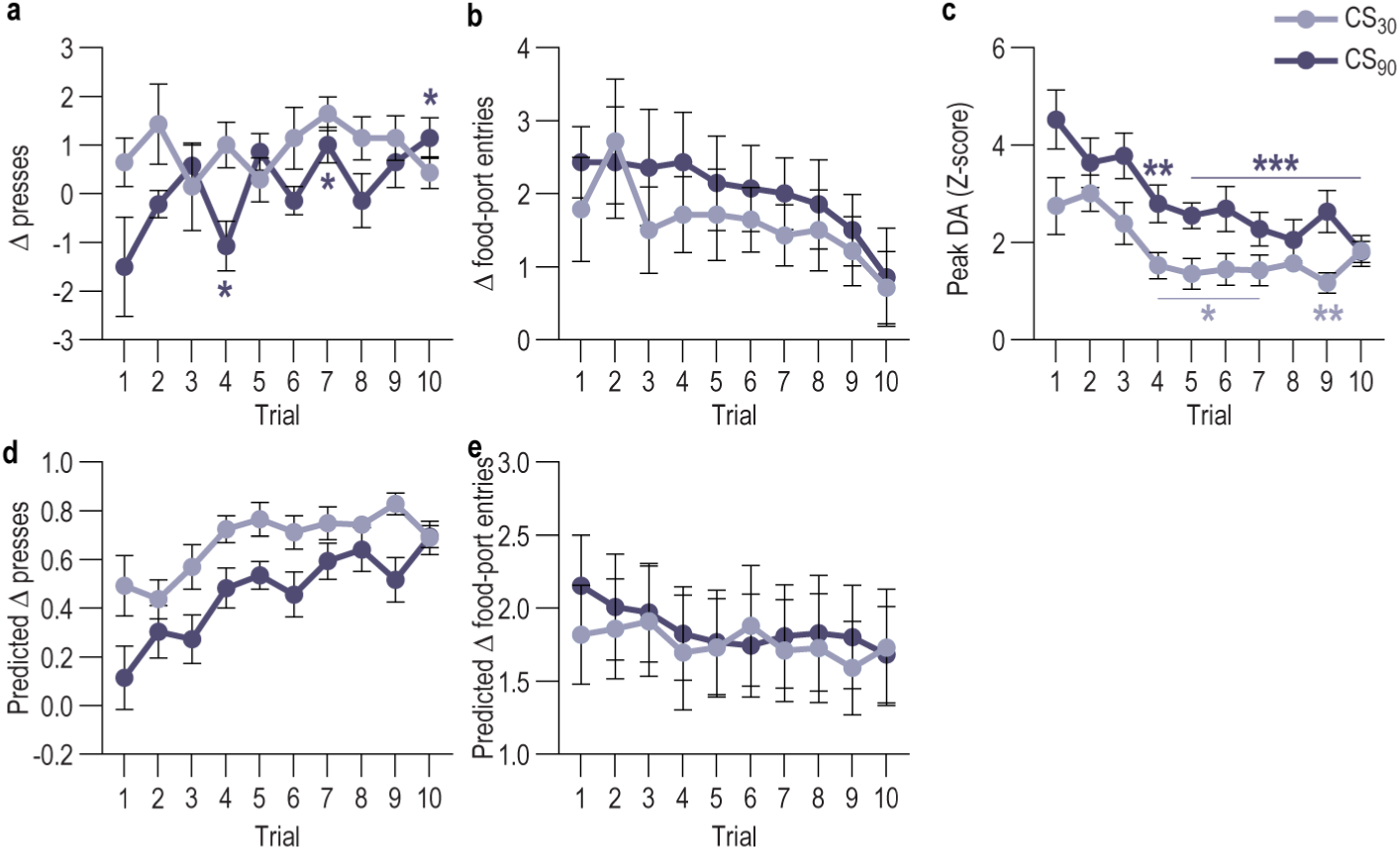
Instrumental reward-seeking and -checking behaviors across trials during Pavlovian-to-instrumental transfer test and seeking and checking behaviors predicted by cue-evoked dopamine. ***A-B,*** Change (Δ) in lever presses *(A)* and food-port entries, excluding those directly linked to lever presses *(B)*, during the cue (conditioned stimulus, CS) from baseline across trials within test. 2-way RM-ANOVA, Presses, Trial: F_9,117_ = 2.20, *P* = 0.03; CS type: F_1,13_ = 4.83, *P* = 0.046; Trial x CS type: F_9,117_ = 1.69, *P* = 0.10; Entries, Trial: F_9,117_ = 1.76, *P* = 0.08; CS type: F_1,13_ = 3.17, *P* = 0.09; Trial x CS type: F_9,117_ = 0.22, *P* = 0.99. The high-probability cue initially suppressed instrumental reward seeking, but as the reward prediction weakened with repeated presentation of this cue without reward, came to elicit an instrumental reward-seeking response similar to the low-probability cue. ***C,*** Peak cue-evoked GRAB_DA_ Z-score compared to baseline across trials within test. Mixed-effects analysis, Trial: F_9,117_ = 10.22, *P* < 0.0001; CS type: F_1,13_ = 26.25, *P* = 0.0002; Trial x CS type: F_9,101_ = 1.37, *P* = 0.21. Dopamine responses were larger to the high-probability cue than the low-probability cue. Cue-evoked dopamine declined as the reward prediction weakened across test trials. ***D,*** Cue-induced change in lever presses predicted by cue-evoked dopamine. We used a linear mixed-model analysis to determine the extent to which cue-evoked instrumental reward seeking could be predicted from the cue-evoked dopamine response. To capture both between-subject variability and with-subject variability occurring across the test we used each trial for each subject. The dependent variable was change in lever presses, CS and trial were repeated factors, peak cue-evoked dopamine was the predictor, and we included subject as a random factor to control for incidental variability due to expression density and fiber placement. β = −0.21, standard error (SE) = 0.07, t_136.60_ = −3.06, *P* = 0.003, 95% CI −0.35 - −0.08. ***E,*** Cue-induced change in food-port entries predicted by cue-evoked dopamine. We used a similar linear mixed-model analysis to determine the extent to which cue-evoked conditional reward-checking behavior could be predicted from the cue-evoked dopamine response. The dependent variable was change in food-port entries (excluding those directly linked to lever presses), CS and trial were repeated factors, peak Cue-evoked dopamine was the predictor, and subject was a random factor. β= 0.15, SE = 0.08, t_139.73_ = 1.89, *P* = 0.06, 95% CI −0.007 - 0.30. We were able to reliably, inversely predict cue-induced changes in instrumental reward-seeking behavior from the cue-evoked dopamine response. Thus, between-subjects and across trials, larger cue-evoked NAc core dopamine predicts a lower instrumental reward-seeking response. Data presented as mean ± s.e.m. \**P* < 0.05, \*\**P* < 0.01, Bonferroni corrected, relative to the first trial.

**Extended Data Figure 1-5.**
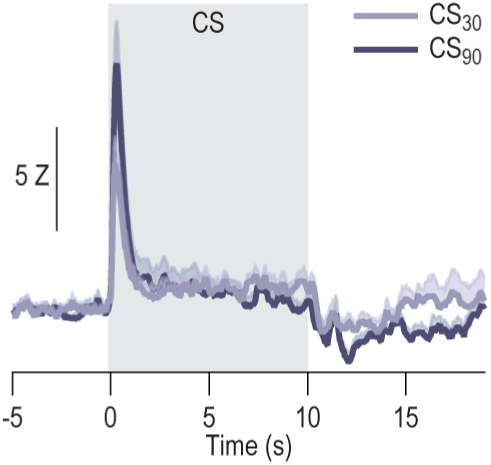
Cue-evoked NAc dopamine without 415 nm signal subtraction. GRAB_DA_ fluorescence changes (drift-corrected 470 nm without subtraction of 415 nm signal) in response to CS presentation during test. Data presented as mean ± s.e.m. As shown in the main data, the onset of both cues triggered robust, transient dopamine release that was larger for the high-probability cue than the low-probability cue.

**Extended Data Figure 1-6.**
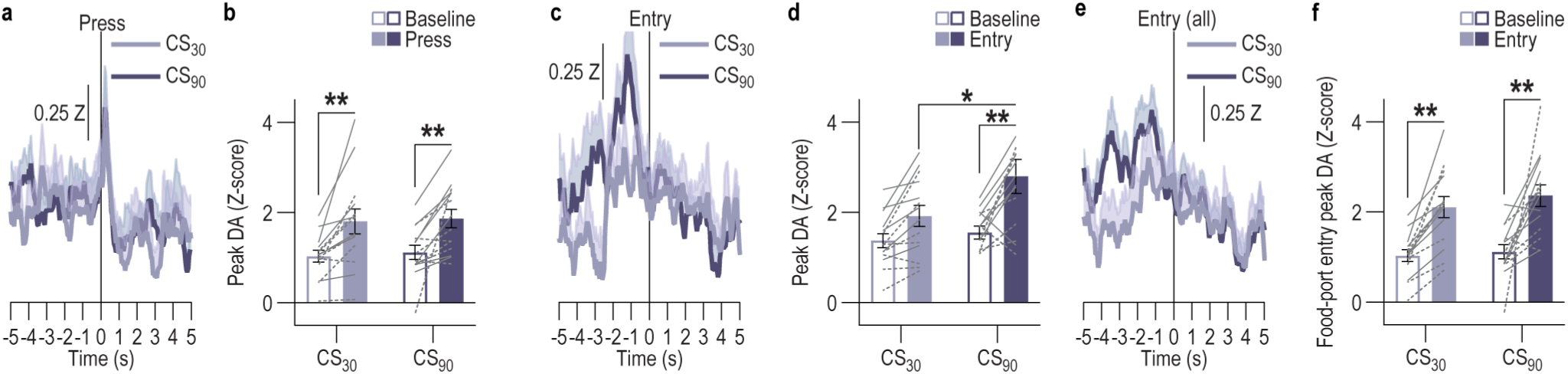
NAc dopamine around cue-induced reward seeking and checking behaviors. ***A,*** GRAB_DA_ fluorescence changes (Z-score) around lever presses during the CSs. ***B,*** Peak GRAB_DA_ Z-score before press during CS. 2-way RM-ANOVA, Press v. baseline: F_1,13_ = 24.22, *P* = 0.0003; CS type: F_1,13_ = 0.14, *P* = 0.71; Press x CS type: F_1,13_ = 0.007, *P* = 0.94. ***C,*** GRAB_DA_ fluorescence changes (Z-score) around food-port entries, excluding those directly linked to lever presses, during the CSs. ***D,*** Peak GRAB_DA_ Z-score before food-port entry. 2-way RM-ANOVA, Entry v. baseline: F_1,13_ = 14.48, *P* = 0.002; CS type: F_1,13_ = 7.49, *P* = 0.02; Entry x CS type: F_1,13_ = 1.99, *P* = 0.18. ***E,*** GRAB_DA_ fluorescence changes (Z-score) around all food-port entries during CS at test. ***F,*** Peak GRAB_DA_ Z-score within 4 s before food-port entry. 2-way RM-ANOVA, Entry v. baseline: F_1,13_ = 48.26, *P* < 0.0001; CS type: F_1,13_ = 1.09, *P* = 0.31; Entry x CS type: F_1,13_ = 0.19, *P* = 0.67. *N* = 14 rats (7 male). There is a significant elevation in dopamine prior to all entries into the food-delivery port (whether linked or not to presses) during both cues. *N* = 14 rats (7 male). Data presented as mean ± s.e.m. Males = solid lines, Females = dashed lines. \**P* < 0.05, \*\**P* < 0.01, \*\*\**P* < 0.001, Bonferroni corrected.

**Extended Data Figure 1-7.**
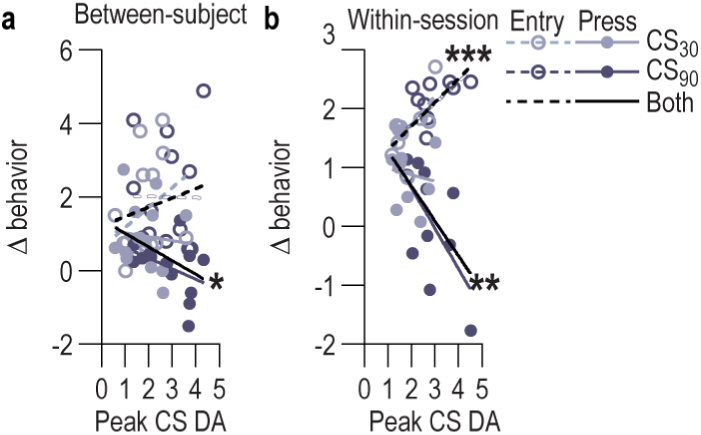
Relationship between cue-evoked dopamine and instrumental reward-seeking and reward-checking behavior. ***A,*** Between-subject (averaged across trial for each subject) correlation between peak cue-evoked GRAB_DA_ Z-score and change in lever presses and food-port entries, excluding those directly linked to lever presses. 2-tailed Pearson correlation, Presses, r_28_ = −0.42, *P* = 0.03, Entries, r_28_ = 0.19, *P* = 0.34. Between subjects there is a negative relationship between cue-evoked dopamine and cue-evoked instrumental reward seeking. Subjects for which the cues elicited a larger dopamine response, on average, showed a lower instrumental reward-seeking response. ***B,*** Within-session (averaged across subjects for each trial) correlation between peak cue-evoked GRAB_DA_ Z-score and change in lever presses and food-port entries, excluding those directly linked to lever presses, for the 15-s CS period. 2-tailed Pearson correlation, Presses, r_20_ = −0.61, *P* = 0.004, Entries, r_20_ = 0.69, *P* = 0.0009. Across trials, cue-evoked dopamine positively relates to cue-induced reward checking and negatively relates to cue-induced instrumental reward seeking. On trials in which the cue elicited a larger dopamine response it also tended to trigger a larger reward-checking and smaller instrumental reward-seeking response. \**P* < 0.05, \*\**P* < 0.01, \*\*\**P* < 0.001.

**Extended Data Figure 2-1.**
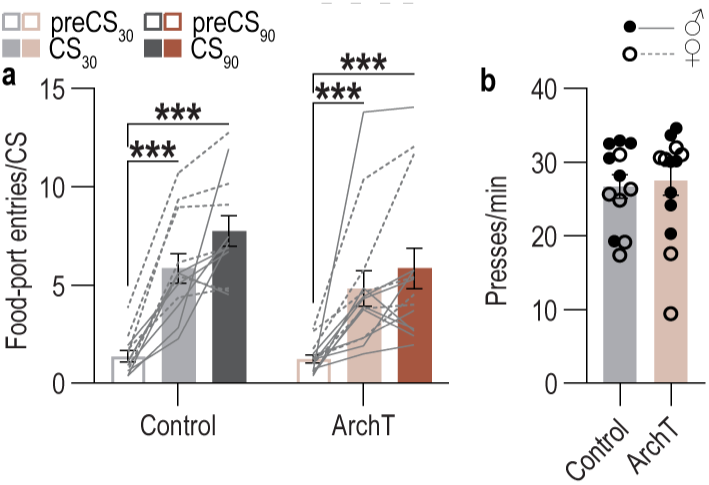
Subjects in the VTA_DA_→NAc inhibition experiment learned the Pavlovian reward-probability predictions and acquire instrumental behavior. ***A,*** Food-port entries per 10-s pre-CS baseline and CS period averaged across the last 2 days of Pavlovian training. 2-way RM-ANOVA, CS period: F_2,48_ = 53.48, *P* < 0.0001; Virus: F_1,24_ = 1.48, *P* = 0.24; Virus x CS period: F_2,48_ = 1.28, *P* = 0.29. Both groups of rats acquired a conditional goal-approach response and learned the differential reward-probability predictions. By the end of training, they showed stronger conditional checks of the food-delivery port to the high-probability cue than the low-probability cue. ***B,*** Lever presses averaged across the last 2 days of instrumental training. 2-tailed t-test, t_24_ = 0.30, *P* = 0.77, 95% CI −4.57 to 6.12. Rats in both groups similarly acquired the instrumental behavior. Control, *N* = 12 rats (6 male; 9 Th-cre+, 3 TH-cre-) ArchT, *N* = 14 Th-cre+ rats (8 male). Data presented as mean ± s.e.m. Males = close circles/solid lines, Females = open circles/dashed lines. \**P* < 0.05, \*\*\**P* < 0.001, Bonferroni corrected.

**Extended Data Figure 2-2.**
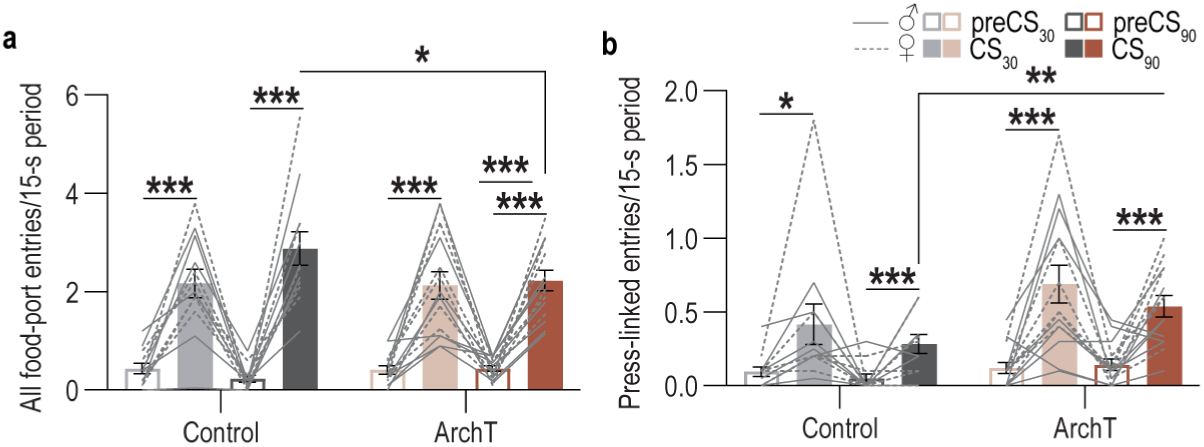
Effect of VTA_DA_→NAc inhibition on all and press-linked cue-evoked entries into the food-delivery port during Pavlovian-to-instrumental transfer. ***A,*** All food-port entries for the 15-s CS (10-s CS + 5-s postCS) and preceding baseline period. 3-way RM-ANOVA, Virus x CS presence x CS type: F_1,24_ = 2.76, *P* = 0.11; CS presence x CS type: F_1,24_ = 3.70, *P* = 0.07; CS presence: F_1,24_ = 166.40, *P* < 0.0001; Virus: F_1,24_ = 0.54, *P* = 0.47; CS type: F_1,24_ = 1.52, *P* = 0.23; Virus x CS presence: F_1,24_ = 2.04, *P* = 0.17; Virus x CS type: F_1,24_ = 0.54, *P* = 0.47. Inhibition of cue-evoked NAc dopamine release tended to reduce overall entries into the food-delivery port, similar to the effect seen only cue-evoked entries that were not directly linked to lever pressing. ***B,*** Press-linked food-port entries (entry occurring directly after and within 2-s of a press) for the 15-s CS and preceding baseline periods. 3-way RM-ANOVA, Virus x CS presence: F_1,24_ = 3.80, *P* = 0.06; Virus: F_1,24_ = 5.98, *P* = 0.02; CS presence: F_1,24_ = 51.09, *P* < 0.0001; CS presence x CS type: F_1,24_ = 1.61, *P* = 0.22; CS type: F_1,24_ = 2.31, *P* = 0.14; Virus x CS type: F_1,24_ = 0.06, *P* = 0.81; Virus x CS presence x CS type: F_1,24_ = 0.17, *P* = 0.69. Inhibition of cue-evoked NAc dopamine release tended to increase entries into the food-delivery port that are directly linked to lever presses, similar to the effect seen lever pressing itself. Control, *N* = 12 rats (6 male; 9 Th-cre+, 3 TH-cre-) ArchT, *N* = 14 Th-cre+ rats (8 male). Data presented as mean ± s.e.m. Males = solid lines, Females = dashed lines. \**P* < 0.05, \*\**P* < 0.01, \*\*\**P* < 0.001, Bonferroni corrected.

**Extended Data Figure 2-3.**
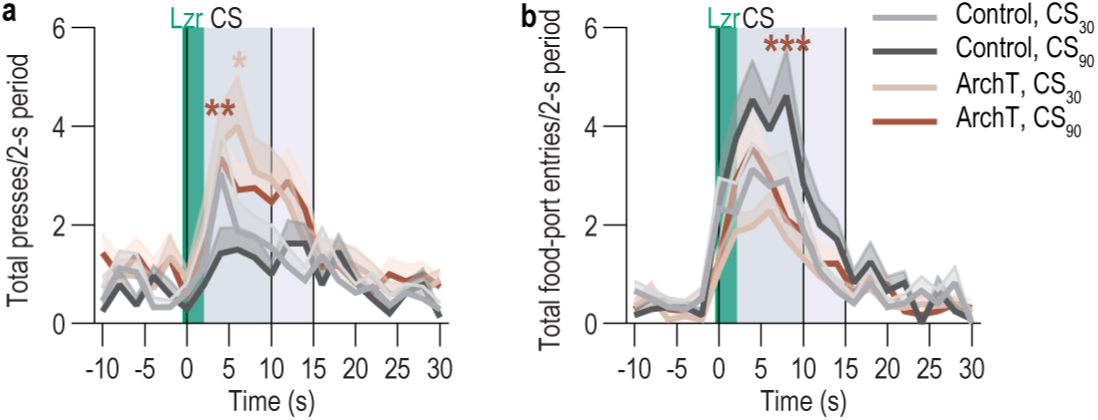
The transient cue-evoked NAc core dopamine response influences behavior throughout the cue. ***A-B,*** Lever presses *(A)* and food-port entries, excluding those directly linked to lever presses, *(B)* in 2-s bins before, during, and after cue (conditioned stimulus, CS) presentation. Gray, 10-s CS; Light gray, 5-s post-CS period; Green, 2.5-s laser (lzr) period. 3-way RM-ANOVA, Presses, Virus x Time: F_20,480_ = 1.64, *P* = 0.04; Time x CS type: F_20,480_ = 1.75, *P* = 0.02; Virus: F_1,24_ = 5.27, *P* = 0.03; Time: F_20,480_ = 12.09, *P* < 0.0001; CS type: F_1,24_ = 0.47, *P* = 0.50; Virus x CS type: F_1,24_ = 0.16, *P* = 0.69; Virus x Time x CS type: F_20,480_ = 0.79, *P* = 0.72; Entries, Virus x Time: F_20,480_ = 2.15, *P* = 0.003; Time x CS type: F_20,480_ = 3.14, *P* < 0.0001; Virus: F_1,24_ = 5.47, *P* = 0.03; Time: F_20,480_ = 35.76, *P* < 0.0001; CS type: F_1,24_ = 14.50, *P* = 0.0009; Virus x CS type: F_1,24_ = 0.28, *P* = 0.60; Virus x Time x CS type: F_20,480_ = 0.70, *P* = 084. Although inhibition of VTA_DA_→NAc projections was restricted to the first 2 s of the cue, this was sufficient to cause a sustained effect to elevate instrumental reward seeking (lever presses) and decrease reward checking (food-port entries) throughout the duration of the cue and post-cue period. Control, *N* = 12 rats (6 male; 9 Th-cre+, 3 TH-cre-) ArchT, *N* = 14 Th-cre+ rats (8 male). Data presented as mean ± s.e.m. Males = solid lines, Females = dashed lines. \**P* < 0.05, \*\**P* < 0.01 ArchT relative to control, Bonferroni corrected.

## REFERENCES

Aitken TJ, Greenfield VY, Wassum KM (2015) Nucleus accumbens core dopamine signaling tracks the need-based motivational value of food-paired cues. J Neurochem.

Aitken TJ, Greenfield VY, Wassum KM (2016) Nucleus accumbens core dopamine signaling tracks the need-based motivational value of food-paired cues. J Neurochem 136:1026–1036.

Anselme P, Güntürkün O (2018) How foraging works: uncertainty magnifies food-seeking motivation. Behav Brain Sci:1–106.

Azrin NH, Hake DF (1969) Positive conditioned suppression: conditioned suppression using positive reinforcers as the unconditioned stimuli. J Exp Anal Behav 12:167–173.

Balleine BW, Ostlund SB (2007) Still at the choice-point: action selection and initiation in instrumental conditioning. Ann N Y Acad Sci 1104:147–171.

Beeler JA, Frazier CR, Zhuang X (2012) Putting desire on a budget: dopamine and energy expenditure, reconciling reward and resources. Front Integr Neurosci 6:49.

Berke JD (2018) What does dopamine mean? Nature Neuroscience 21:787–793.

Bernklau TW, Righetti B, Mehrke LS, Jacob SN (2024) Striatal dopamine signals reflect perceived cue-action-outcome associations in mice. Nat Neurosci 27:747–757.

Bornhoft KN, Prohofsky J, O’Neal TJ, Wolff AR, Saunders BT (2025) Striatal dopamine represents valence on dynamic regional scales. J Neurosci 45.

Collins AL, Aitken TJ, Greenfield VY, Ostlund SB, Wassum KM (2016) Nucleus Accumbens Acetylcholine Receptors Modulate Dopamine and Motivation. Neuropsychopharmacology.

Collins AL, Aitken TJ, Huang IW, Shieh C, Greenfield VY, Monbouquette HG, Ostlund SB, Wassum KM (2019) Nucleus Accumbens Cholinergic Interneurons Oppose Cue-Motivated Behavior. Biol Psychiatry.

Colwill RM, Rescorla RA (1990) Effect of reinforcer devaluation on discriminative control of instrumental behavior. J Exp Psychol Anim Behav Process 16:40–47.

Corbit LH, Balleine BW (2011) The general and outcome-specific forms of Pavlovian-instrumental transfer are differentially mediated by the nucleus accumbens core and shell. J Neurosci 31:11786–11794.

Corbit LH, Balleine BW (2016) Learning and Motivational Processes Contributing to Pavlovian-Instrumental Transfer and Their Neural Bases: Dopamine and Beyond. Curr Top Behav Neurosci.

Crombag HS, Galarce EM, Holland PC (2008) Pavlovian influences on goal-directed behavior in mice: the role of cue-reinforcer relations. Learn Mem 15:299–303.

Day JJ, Roitman MF, Wightman RM, Carelli RM (2007) Associative learning mediates dynamic shifts in dopamine signaling in the nucleus accumbens. Nat Neurosci 10:1020–1028.

Day JJ, Jones JL, Wightman RM, Carelli RM (2010) Phasic nucleus accumbens dopamine release encodes effort- and delay-related costs. Biol Psychiatry 68:306–309.

du Hoffmann J, Nicola SM (2014) Dopamine invigorates reward seeking by promoting cue-evoked excitation in the nucleus accumbens. J Neurosci 34:14349–14364.

Eshel N, Tian J, Bukwich M, Uchida N (2016) Dopamine neurons share common response function for reward prediction error. Nat Neurosci 19:479–486.

Fiorillo CD, Tobler PN, Schultz W (2003) Discrete coding of reward probability and uncertainty by dopamine neurons. Science 299:1898–1902.

Flagel SB, Clark JJ, Robinson TE, Mayo L, Czuh A, Willuhn I, Akers CA, Clinton SM, Phillips PEM, Akil H (2011) A selective role for dopamine in stimulus-reward learning. Nature 469:53–57.

Floresco SB, Magyar O, Ghods-Sharifi S, Vexelman C, Tse MT (2006) Multiple dopamine receptor subtypes in the medial prefrontal cortex of the rat regulate set-shifting. Neuropsychopharmacology 31:297–309.

Gan JO, Walton ME, Phillips PE (2009) Dissociable cost and benefit encoding of future rewards by mesolimbic dopamine. Nat Neurosci 13:25–27.

Garr E, Cheng Y, Jeong H, Brooke S, Castell L, Bal A, Magnard R, Namboodiri VMK, Janak PH (2024) Mesostriatal dopamine is sensitive to changes in specific cue-reward contingencies. Sci Adv 10:eadn4203.

Glimcher PW (2011) Understanding dopamine and reinforcement learning: the dopamine reward prediction error hypothesis. Proc Natl Acad Sci U S A 108 Suppl 3:15647–15654.

Greenstreet F, Vergara HM, Johansson Y, Pati S, Schwarz L, Lenzi SC, Geerts JP, Wisdom M, Gubanova A, Rollik LB, Kaur J, Moskovitz T, Cohen J, Thompson E, Margrie TW, Clopath C, Stephenson-Jones M (2025) Dopaminergic action prediction errors serve as a value-free teaching signal. Nature 643:1333–1342.

Halbout B, Hutson C, Wassum KM, Ostlund SB (2022) Dorsomedial prefrontal cortex activation disrupts Pavlovian incentive motivation. Front Behav Neurosci 16:999320.

Halbout B, Marshall AT, Azimi A, Liljeholm M, Mahler SV, Wassum KM, Ostlund SB (2019) Mesolimbic dopamine projections mediate cue-motivated reward seeking but not reward retrieval in rats. Elife 8.

Hamid AA, Pettibone JR, Mabrouk OS, Hetrick VL, Schmidt R, Vander Weele CM, Kennedy RT, Aragona BJ, Berke JD (2015) Mesolimbic dopamine signals the value of work. Nat Neurosci.

Hart AS, Clark JJ, Phillips PE (2015) Dynamic shaping of dopamine signals during probabilistic Pavlovian conditioning. Neurobiol Learn Mem 117:84–92.

Hart AS, Rutledge RB, Glimcher PW, Phillips PE (2014) Phasic dopamine release in the rat nucleus accumbens symmetrically encodes a reward prediction error term. J Neurosci 34:698–704.

Holland PC (2004) Relations between Pavlovian-instrumental transfer and reinforcer devaluation. J Exp Psychol Anim Behav Process 30:104–117.

Howe MW, Dombeck DA (2016) Rapid signalling in distinct dopaminergic axons during locomotion and reward. Nature 535:505–510.

Jean-Richard-Dit-Bressel P, Clifford CWG, McNally GP (2020) Analyzing Event-Related Transients: Confidence Intervals, Permutation Tests, and Consecutive Thresholds. Front Mol Neurosci 13:14.

Jentsch JD, Pennington ZT (2014) Reward, interrupted: Inhibitory control and its relevance to addictions. Neuropharmacology 76 Pt B:479–486.

Kahnt T, Schoenbaum G (2025) The curious case of dopaminergic prediction errors and learning associative information beyond value. Nat Rev Neurosci 26:169–178.

Kalmbach A, Winiger V, Jeong N, Asok A, Gallistel CR, Balsam PD, Simpson EH (2022) Dopamine encodes real-time reward availability and transitions between reward availability states on different timescales. Nat Commun 13:3805.

Kaźmierczak M, Nicola SM (2022) The Arousal-motor Hypothesis of Dopamine Function: Evidence that Dopamine Facilitates Reward Seeking in Part by Maintaining Arousal. Neuroscience 499:64–103.

Konorski J (1967) Integrative activity of the brain: An interdisciplinary approach. Chicago: University of Chicago Press.

Kutlu MG, Zachry JE, Melugin PR, Cajigas SA, Chevee MF, Kelly SJ, Kutlu B, Tian L, Siciliano CA, Calipari ES (2021) Dopamine release in the nucleus accumbens core signals perceived saliency. Curr Biol 31:4748–4761.e4748.

Kutlu MG, Zachry JE, Melugin PR, Tat J, Cajigas S, Isiktas AU, Patel DD, Siciliano CA, Schoenbaum G, Sharpe MJ, Calipari ES (2022) Dopamine signaling in the nucleus accumbens core mediates latent inhibition. Nat Neurosci 25:1071–1081.

Le Heron C, Apps MAJ, Husain M (2018) The anatomy of apathy: A neurocognitive framework for amotivated behaviour. Neuropsychologia 118:54–67.

Lerner TN, Holloway AL, Seiler JL (2021) Dopamine, Updated: Reward Prediction Error and Beyond. Curr Opin Neurobiol 67:123–130.

Levy R, Dubois B (2006) Apathy and the functional anatomy of the prefrontal cortex-basal ganglia circuits. Cereb Cortex 16:916–928.

Lex A, Hauber W (2008) Dopamine D1 and D2 receptors in the nucleus accumbens core and shell mediate Pavlovian-instrumental transfer. Learn Mem 15:483–491.

Lichtenberg NT, Wassum KM (2016) Amygdala mu-opioid receptors mediate the motivating influence of cue-triggered reward expectations. Eur J Neurosci.

Lichtenberg NT, Pennington ZT, Holley SM, Greenfield VY, Cepeda C, Levine MS, Wassum KM (2017) Basolateral amygdala to orbitofrontal cortex projections enable cue-triggered reward expectations. J Neurosci.

Lopes G, Bonacchi N, Frazão J, Neto JP, Atallah BV, Soares S, Moreira L, Matias S, Itskov PM, Correia PA, Medina RE, Calcaterra L, Dreosti E, Paton JJ, Kampff AR (2015) Bonsai: an event-based framework for processing and controlling data streams. Front Neuroinform 9:7.

Lovibond PF (1981) Appetitive Pavlovian-instrumental interactions: effects of inter-stimulus interval and baseline reinforcement conditions. Q J Exp Psychol B 33:257–269.

Maes EJP, Sharpe MJ, Usypchuk AA, Lozzi M, Chang CY, Gardner MPH, Schoenbaum G, Iordanova MD (2020) Causal evidence supporting the proposal that dopamine transients function as temporal difference prediction errors. Nat Neurosci 23:176–178.

Malvaez M, Greenfield VY, Wang AS, Yorita AM, Feng L, Linker KE, Monbouquette HG, Wassum KM (2015) Basolateral amygdala rapid glutamate release encodes an outcome-specific representation vital for reward-predictive cues to selectively invigorate reward-seeking actions. Sci Rep 5:12511.

Marshall AT, Ostlund SB (2018) Repeated cocaine exposure dysregulates cognitive control over cue-evoked reward-seeking behavior during Pavlovian-to-instrumental transfer. Learn Mem 25:399–409.

Marshall AT, Munson CN, Maidment NT, Ostlund SB (2020) Reward-predictive cues elicit excessive reward seeking in adolescent rats. Dev Cogn Neurosci 45:100838.

Marshall AT, Halbout B, Munson CN, Hutson C, Ostlund SB (2023) Flexible control of Pavlovian-instrumental transfer based on expected reward value. J Exp Psychol Anim Learn Cogn 49:14–30.

Niv Y, Daw ND, Joel D, Dayan P (2007) Tonic dopamine: opportunity costs and the control of response vigor. Psychopharmacology (Berl) 191:507–520.

Ostlund S, Marshall A (2021) Probing the role of reward expectancy in Pavlovian-instrumental transfer. Current Opinion in Behavioral Sciences 41:106–113.

Ostlund SB, Wassum KM, Murphy NP, Balleine BW, Maidment NT (2011) Extracellular Dopamine Levels in Striatal Subregions Track Shifts in Motivation and Response Cost during Instrumental Conditioning. Journal of Neuroscience 31:200–207.

Parker NF, Cameron CM, Taliaferro JP, Lee J, Choi JY, Davidson TJ, Daw ND, Witten IB (2016) Reward and choice encoding in terminals of midbrain dopamine neurons depends on striatal target. Nat Neurosci 19:845–854.

Rescorla R (1992) Response-outcome versus outcome-response associations in instrumental learning. Animal Learning & Behavior 20:223–232.

Rescorla RA (1994) Transfer of instrumental control mediated by a devalued outcome. Animal Learning & Behavior 22:27–33.

Robinson TE, Berridge KC (2025) The Incentive-Sensitization Theory of Addiction 30 Years On. Annu Rev Psychol 76:29–58.

Roitman MF, Stuber GD, Phillips PE, Wightman RM, Carelli RM (2004) Dopamine operates as a subsecond modulator of food seeking. J Neurosci 24:1265–1271.

Saddoris MP, Sugam JA, Stuber GD, Witten IB, Deisseroth K, Carelli RM (2015) Mesolimbic dopamine dynamically tracks, and is causally linked to, discrete aspects of value-based decision making. Biol Psychiatry 77:903–911.

Saunders BT, Robinson TE (2012) The role of dopamine in the accumbens core in the expression of Pavlovian-conditioned responses. Eur J Neurosci 36:2521–2532.

Schultz W (2002) Getting formal with dopamine and reward. Neuron 36:241–263.

Schultz W (2007) Multiple dopamine functions at different time courses. Annu Rev Neurosci 30:259–288.

Schultz W (2016) Dopamine reward prediction-error signalling: a two-component response. Nat Rev Neurosci 17:183–195.

Schultz W, Dayan P, Montague PR (1997) A neural substrate of prediction and reward. Science 275:1593–1599.

Sias A, Morse A, Wang S, Greenfield V, Goodpaster C, Wrenn T, Wikenheiser A, Holley S, Cepeda C, Levine M, Wassum K (2021) A bidirectional corticoamygdala circuit for the encoding and retrieval of detailed reward memories. eLife 10.

Sias AC, Jafar Y, Goodpaster CM, Ramírez-Armenta K, Wrenn TM, Griffin NK, Patel K, Lamparelli AC, Sharpe MJ, Wassum KM (2024) Dopamine projections to the basolateral amygdala drive the encoding of identity-specific reward memories. Nat Neurosci 27:728–736.

Steinberg EE, Keiflin R, Boivin JR, Witten IB, Deisseroth K, Janak PH (2013) A causal link between prediction errors, dopamine neurons and learning. Nat Neurosci 16:966–973.

Stephens DW, Krebs JR (1986) Foraging theory: Princeton university press.

Sun F et al. (2018) A Genetically Encoded Fluorescent Sensor Enables Rapid and Specific Detection of Dopamine in Flies, Fish, and Mice. Cell 174:481–496.e419.

Syed EC, Grima LL, Magill PJ, Bogacz R, Brown P, Walton ME (2016) Action initiation shapes mesolimbic dopamine encoding of future rewards. Nat Neurosci 19:34–36.

Taira M, Millard SJ, Verghese A, DiFazio LE, Hoang IB, Jia R, Sias A, Wikenheiser A, Sharpe MJ (2024) Dopamine Release in the Nucleus Accumbens Core Encodes the General Excitatory Components of Learning. J Neurosci 44.

Timberlake W (1994) Behavior systems, associationism, and Pavlovian conditioning. Psychon Bull Rev 1:405–420.

Tobler PN, Fiorillo CD, Schultz W (2005) Adaptive coding of reward value by dopamine neurons. Science 307:1642–1645.

Waelti P, Dickinson A, Schultz W (2001) Dopamine responses comply with basic assumptions of formal learning theory. Nature 412:43–48.

Wassum KM, Ostlund SB, Loewinger GC, Maidment NT (2013) Phasic Mesolimbic Dopamine Release Tracks Reward Seeking During Expression of Pavlovian-to-Instrumental Transfer. Biol Psychiatry 73:747–755.

Watson P, Wiers RW, Hommel B, de Wit S (2018) Motivational sensitivity of outcome-response priming: Experimental research and theoretical models. Psychon Bull Rev 25:2069–2082.

